# Stay or Stray: Lpar1 regulates neutrophil retention and epidermal homeostasis in early zebrafish development

**DOI:** 10.1101/2025.06.22.660980

**Authors:** Shih-Chi Li, Yu-Chi Lin, Chung-Der Hsiao, Shyh-Jye Lee

**Affiliations:** Department of Life Science, National Taiwan University, 1 Roosevelt Rd., Sec., 4, Taipei 10617, Taiwan, R.O.C; Department of Bioscience Technology, Chung Yuan Christian University, Chung-Li, Taiwan 32023, R.O.C; Research Center for Developmental Biology and Regenerative, National Taiwan University, 1 Roosevelt Rd., Sec., 4, Taipei 10617, Taiwan, R.O.C; Center for Biotechnology, National Taiwan University, 1 Roosevelt Rd., Sec., 4, Taipei 10617, Taiwan, R.O.C

**Keywords:** Zebrafish, lysophosphatidic acid receptor 1 (Lpar1), neutrophils, caudal hematopoietic tissue, chemokine (C-X-C motif) ligand 12 (CXCL12)

## Abstract

Neutrophils are the most abundant myeloid cells in the vertebrate innate immune system and play a crucial role in host defense. Dysregulated release from hematopoietic tissue or excessive recruitment to inflamed sites is associated with immunopathologies, including chronic inflammation and recurrent infections. Lysophosphatidic acid (LPA) is a bioactive lipid that signals through a family of G protein-coupled receptors (LPAR1-6). Among these, LPA-LPAR1 signaling is activated in inflamed tissues and is known to modulate neutrophil infiltration during inflammatory responses. However, its role in neutrophil dynamics during early development remains unclear. Here, we report a novel function of Lpar1 in regulating neutrophil behavior during early zebrafish development. In Lpar1-deficient embryos, we observed a significant increase in neutrophil dispersal from the caudal hematopoietic tissue (CHT) at 3 days post-fertilization (dpf), with most dispersed neutrophils infiltrating the skin and contributing to elevated inflammatory signaling. Prior to dispersal, Lpar1-deficient embryos exhibited increased apoptosis of superficial epidermal cells and reduced expression of *cxcl12a*, a key retention signal for neutrophils in the CHT. Together, these findings highlight an essential role for Lpar1 in maintaining neutrophil retention and epidermal homeostasis during early development, thereby limiting inappropriate inflammation.

**SUMMARY STATEMENT:** This work identifies a new role for a known inflammatory receptor in coordinating immune cell retention and skin stability during early development.

## INTRODUCTION

The inappropriate release of innate immune cells from hematopoietic tissue is associated with immunodeficiency diseases such as sepsis and WHIM syndrome (Kawai & Malech, 2009; Shen et al., 2021; van der Poll et al., 2021). Excessive release and recruitment of immune cells can lead to chronic inflammation, while impaired mobilization of these cells contributes to recurrent infections. Neutrophils, the most abundant myeloid cells in peripheral blood, first respond to infection or tissue damage. In adult mammals, neutrophils primarily develop in the bone marrow and are released into the bloodstream upon maturation. The bone marrow maintains the homeostasis of circulating neutrophils by releasing the chemokine CXCL12 (SDF-1), which interacts with its receptor CXCR4 on neutrophils to regulate their retention and release (Link, 2005; Suratt et al., 2004). In response to bacterial infection or tissue damage, neutrophils are recruited from the bloodstream to inflamed sites by detecting various molecules, such as danger-associated molecular patterns (DAMPs), hydrogen peroxide (H₂O₂), lipids, and chemokines (de Oliveira et al., 2016). In addition, local inflammation can trigger the release of neutrophils from the bone marrow via inflammatory mediators (Burdon et al., 2005; Martin et al., 2003). The resolution of infiltrated neutrophils is crucial to prevent their prolonged activation in inflamed tissue. Typically, neutrophils undergo apoptosis or necrosis followed by macrophage efferocytosis or reverse migration back into the bloodstream (Buckley et al., 2013; Mathias et al., 2006; Woodfin et al., 2011). Altogether, understanding the molecular networks that regulate the release, recruitment, and resolution of neutrophils during infection or tissue damage is essential and can provide further insights into treating immunodeficiency diseases.

Lysophosphatidic acid (LPA) is a small bioactive phospholipid present in various tissues and fluids, mediating a broad range of cellular and signaling functions, including cell proliferation, cell migration, cell survival, cytokine secretion, and inflammation (Geraldo et al., 2021). LPA is mainly generated extracellularly by the plasma enzyme autotoxin (ATX), which converts lysophospholipids (LPs) into LPA (Aikawa et al., 2015). LPA exerts its extracellular signaling by binding to its six specific G protein-coupled receptors (GPCRs), known as LPA receptors 1-6 (LPAR1-6), with each LPAR activating different intracellular mediators. LPAR1, the first identified LPA receptor, is ubiquitously expressed in various organs such as the brain, heart, intestine, lung, kidney, spleen, and skeletal muscle. (Geraldo et al., 2021; Tager et al., 2008). LPAR1 is strongly expressed in chronic inflammation tissues, such as in *Candida albicans* water-soluble fraction (CAWS)-induced vasculitis and rheumatoid arthritis (RA) (Miyabe et al., 2019; Miyabe et al., 2013; Zhao et al., 2008). Inhibition of LPAR1 signaling has been shown to ameliorate disease progression and reduce immune cell infiltration, including neutrophils. These studies demonstrate that LPA-LPAR1 signaling is essential for the pathogenesis of chronic inflammatory diseases and the modulation of leukocyte migration. However, most research has focused on LPA-LPAR1 signaling in the context of inflammation. The role of LPA-LPAR1 signaling in regulating neutrophil dynamics in healthy conditions or during early development remains unresolved. Zebrafish (*Danio rerio*) have emerged as a great alternative model for studying neutrophil development and dynamics (Harvie & Huttenlocher, 2015). In zebrafish, Lpar1 has been shown to play important roles in early angiogenesis, lymphatic vessel development, and craniofacial chondrogenesis (Lee et al., 2008; Nishioka et al., 2016; Yukiura et al., 2011). Yet, the role of Lpar1 in zebrafish neutrophil dynamics has not yet been examined.

In this study, we used loss-of-function approaches, including MO-mediated knockdown and a genetic mutant strain, to investigate the role of Lpar1 in regulating neutrophil dynamics during zebrafish development. We found that Lpar1-deficient embryos exhibited a significant increase in neutrophil dispersal from hematopoietic tissue, with most neutrophils infiltrating the skin. Delving into the underlying mechanisms, we observed an increase in apoptotic epidermal cells and a downregulation of *cxcl12a*, a key retention signal for neutrophils in hematopoietic tissue, in Lpar1 morphants. Together, these findings reveal a novel role for Lpar1 in maintaining epidermal homeostasis and regulating neutrophil retention, thereby fine-tuning neutrophil dynamics during early zebrafish development.

## MATERIALS & METHODS

### Zebrafish maintenance and strains

Zebrafish (*Danio rerio*) AB strain, *Tg* (*mpx:EGFP*) (Renshaw et al., 2006), *Tg* (*mpeg1:EGFP*) (Ellett et al., 2011), *Tg* (*krt4*: h2afv-mCherry)^cy9^ (Chen et al., 2014), and Lpar1 sa38782 mutant line (Kettleborough et al., 2013) were raised at 28.5℃ with a 14/10-hour light/dark cycle. Embryos were collected by natural spawning and incubated in 0.3 × Danieau’s buffer (1.4 mM NaCl, 210 μM KCl, 120 μM MgSO_4,_ 180 μM Ca(NO_3_), 1.55 mM HEPES in double-distilled water, pH 7.2). Developmental stages of embryos and larvae were defined according to Kimmel et al. (1995). To inhibit pigmentation, embryos were transferred to 0.3X Danieau’s buffer containing 200 μM phenylthiourea (PTU). All experimental procedures adhered to the animal use guidelines established by National Taiwan University, Taipei, Taiwan.

### Ethics Statement

All experimental procedures involving zebrafish were approved by the Laboratory Animal Use Committee of National Taiwan University, Taipei, Taiwan (IACUC Approval ID: NTU-112-EL-00002), and conducted in accordance with the approved guidelines.

### Embryo microinjection

Antisense splice-blocking MO oligonucleotide (MO) targeting the *lpar1* sequence (TGGAGCACTTACCCAATACAATCAC) (Lee et al., 2008) and the standard control MO with a random sequence (std MO: CCTCTTACCTCAGTTACAATTTATA) were synthesized by Gene Tools (Oregon, USA). The Lpar1 expression plasmid, containing the coding region of *lpar1* in a pcDNA3.1/V5-His-TOPO expression vector driven by a cytomegalovirus (CMV) promoter, was previously constructed (Lee et al., 2008).

Plasmid preparations for injection were made using the HiYield™ Plasmid Mini Kit (ARROWTECH, New Taipei City, Taiwan). Cxcl12a RNA for ectopic expression was generated using the GenBuilder Plus Cloning Kit (#L00744, GenScript, New Jersey, USA) by assembling the *cxcl12a* coding region and P2A-mCherry into the pCS2^+^ vector. RNA was synthesized via *in vitro* transcription using the mMESSAGE mMACHINE™ SP6 Transcription Kit (Invitrogen, Cat# AM1340, Waltham, MA, USA). The MO was preheated at 65°C for 10 minutes to prevent precipitation. Injection mixtures were prepared with the MO, plasmid, or RNA dissolved in double-distilled water (ddH2O), with phenol red solution (#RNBG2159, Sigma-Aldrich, St. Louis, Missouri, USA) accounting for 10% of the total volume. Glass capillaries with an outer diameter of 1.14 mm and an inner diameter of 0.5 mm (#4878, World Precision Instruments Inc., Sarasota, FL) were pulled using a horizontal puller (P-97, Sutter Instrument, Novato, CA). A volume of 1 nL of the injection mixture was delivered into zebrafish embryos at the 1-cell stage using an air-pressure-driven ZGB Pco-1500 Plus Liter Microinjector (Zgenebio Biotech Inc., Taipei, Taiwan).

### Genomic DNA collection and genotyping

The genotyping strategy for the Lpar1 sa38782 mutant line is described in Fig. S1. In brief, genomic DNA was extracted from the caudal fin tissue by lysing the tissue and releasing DNA in 100 μL of 50 mM sodium hydroxide (NaOH) at 95°C for 30 minutes using a PCR machine. The pH was then neutralized by adding 1 M Tris-HCl solution (pH 8.0) to prepare the sample for subsequent polymerase chain reaction (PCR). PCR products were digested with AccI (#R0161S, New England Biolabs, Ipswich, Massachusetts, USA) or ClaI (#R0197S, New England Biolabs,) restriction enzymes at 37°C overnight in rCutSmart™ Buffer (provided with the restriction enzymes). After digestion, an equal volume of dilution buffer provided by the Qsep100 capillary electrophoresis analyzer (BiOptic, New Taipei City, Taiwan) was added to the mixtures. Capillary electrophoresis was performed following the manufacturer’s instructions, and the results were analyzed using the manufacturer-provided software, Q-analyzer.

### Immunohistochemistry

Whole-mount immunohistochemistry (IHC) staining was performed as described by Inoue and Wittbrodt (2011), with slight modifications. After blocking, the larvae were incubated with primary antibodies: chick anti-GFP (Aves Labs, 1:500) and rabbit anti-DsRed (Living Colors® DsRed Polyclonal Antibody, Takara Bio, 1:500), and incubated overnight at 4°C. Secondary antibodies, goat anti-chick or anti-rabbit IgG conjugated with Alexa Fluor 488 or Alexa Fluor 568 (Invitrogen, 1:500), were applied, and the samples were incubated at 4°C for 1-2 hours. Following several washes with a blocking buffer, the samples were stored in phosphate-buffered saline (PBS) until image acquisition.

### TUNEL assay

The Click-iT™ Plus TUNEL Alexa Fluor 488 Assay for In Situ Apoptosis Detection (#C10617, Invitrogen, Waltham, Massachusetts, USA) was performed according to the user manual with slight modifications. Larvae at different developmental stages were dechorionated and fixed in freshly prepared 4% paraformaldehyde (PFA) in phosphate-buffered saline (PBS, 137 mM NaCl, 2.7 mM KCl, 10 mM Na_2_HPO_4_, and 1.8 mM KH_2_PO_4_.) overnight at 4°C. For the Click-iT™ Plus reaction, samples were incubated in the Click-iT™ Plus TUNEL reaction cocktail for 30 minutes to 1 hour at 37°C. Following the TUNEL reaction, samples were subjected to IHC procedures, including blocking and primary antibody incubation. After IHC staining, samples were stored in PBS until image acquisition.

### Caspase inhibition

Embryos at 32 hpf were dechorionated and incubated in 100 μM pan-caspase inhibitor zVAD-fmk (#627610, Sigma-Aldrich, St. Louis, MO, USA) prepared in fish water. The treatment continued until 3 dpf, after which embryos were subjected to imaging for neutrophil quantification.

### cDNA preparation and real-time quantitative PCR (RT-qPCR)

Whole larvae were homogenized using a 25-gauge needle and syringe in 600 μL RNAzol® RT solution (Molecular Research Center, Ohio, USA) to extract total RNA. Samples were added with 240 μL diethylpyrocarbonate (DEPC) water, and centrifuged at 14000 RCF for 15 minutes. The aqueous-phase supernatants (500 μL) were concentrated and purified using the RNA Clean & Concentrator™-5 kit (#R1015, Zymo Research, Taipei, Taiwan), following the manufacturer’s instructions. Ethanol (95-100%) was added in equal volume to the supernatants, which were then transferred to Zymo-Spin™ IC Columns for subsequent purification steps according to the manual.

RNA quality was assessed via electrophoresis on a 1% agarose gel in TAE buffer (40mM Tris, 20mM Acetate, and 1mM EDTA) and using the NanoDrop One Spectrophotometer (Thermo Fisher Scientific, Waltham, Massachusetts, USA) for concentration and purity measurements For cDNA synthesis, 2 μg of RNA was annealed with oligo dT primer at 70°C for 5 minutes, followed by reverse transcription using M-MLV Reverse Transcriptase (Promega, Madison, MI, USA) at 37°C for 1 hour. The resulting first-strand cDNA was stored at −20°C for further use.

RT-qPCR was conducted using the CFX 96 Touch Real-Time Detection System (Bio-Rad, Hercules, California, USA) with iQ™ SYBR® Green Supermix (#1708882, Bio-Rad). Specific primers used are listed in Table 1. The thermal cycling conditions involved initial denaturation at 95°C for 2 minutes, followed by 40 cycles of 5 seconds at 95°C and 30 seconds at 60°C. To confirm amplicon specificity, a melting curve analysis was performed by heating from 65°C to 95°C, increasing by 0.5°C increments for 0.05 seconds. Gene expression levels were normalized to the internal control gene, eukaryotic translation elongation factor 1 alpha 1, like 1 (*eef1a1l1*), and relative fold changes for each treatment were calculated using the 2^−ΔΔCt^ method (Livak & Schmittgen, 2001).

**Table 1.**
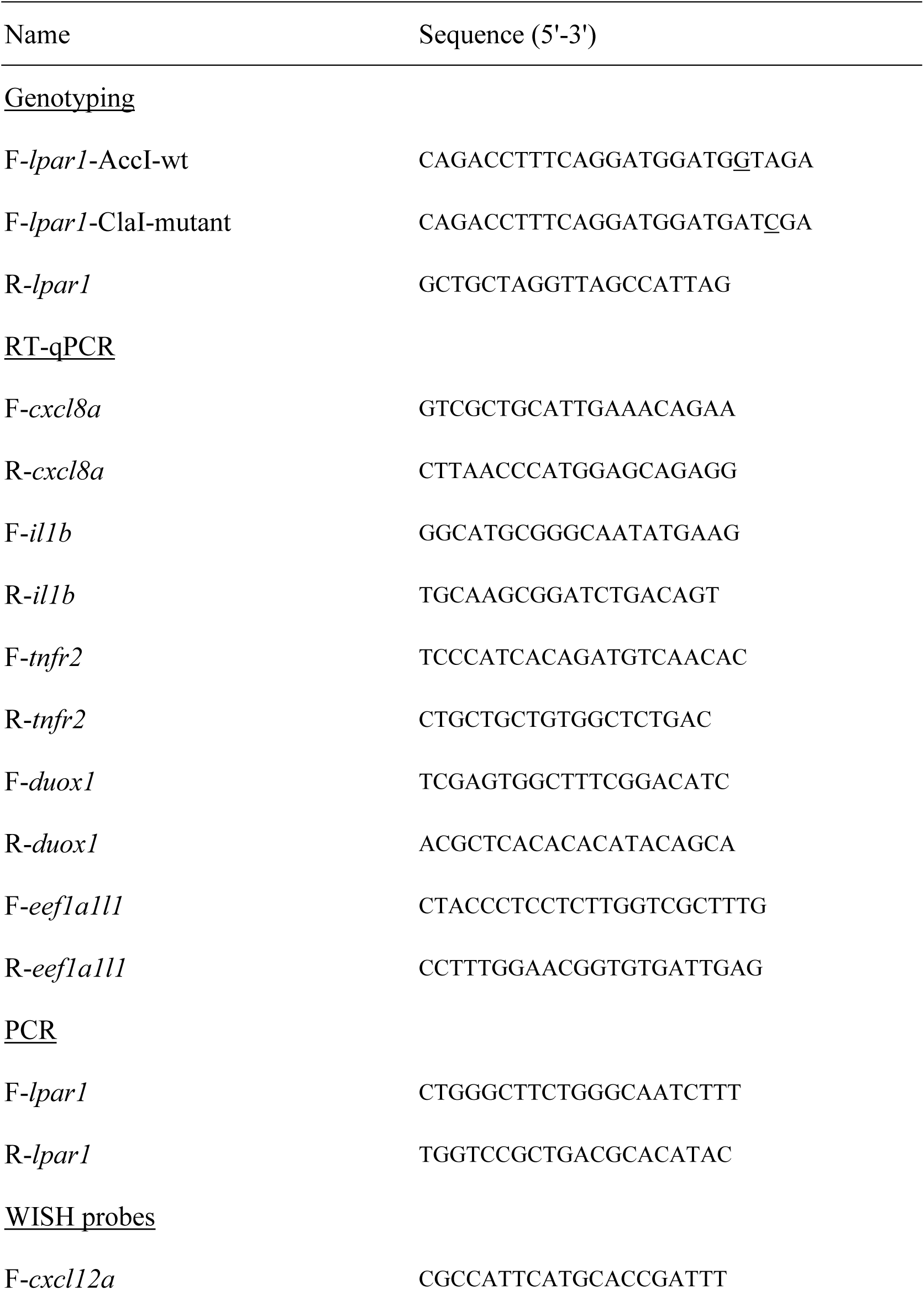
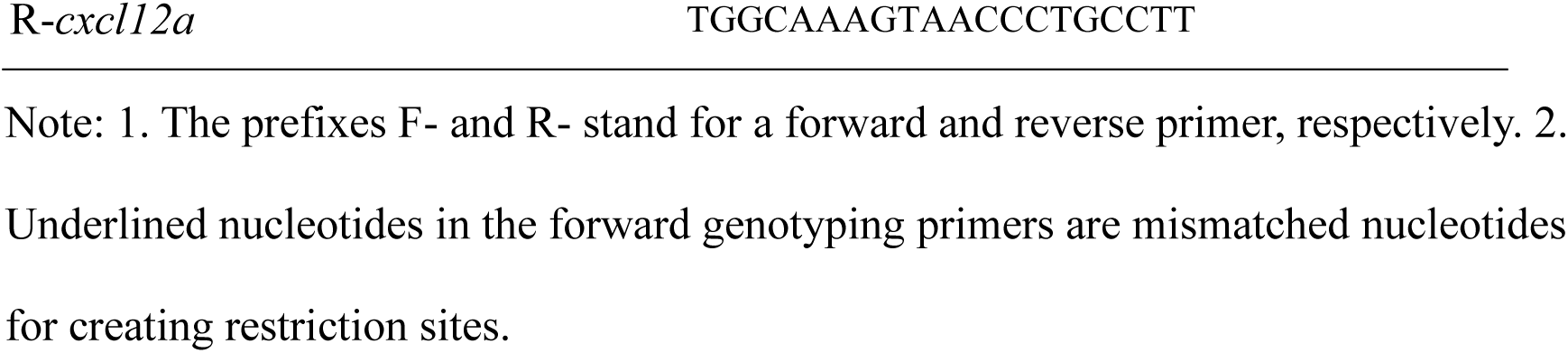
List of genotyping, RT-qPCR, PCR, and WISH probe primers.

### Neutrophil dissociation, ROS detection, and FACS sorting from zebrafish larvae at 3 dp

At 3 dpf, 100 *Tg* (*mpx:EGFP*) larvae from each treatment were dechorionated and pooled for single-cell dissociation. The dissociation was conducted following the method described by Bresciani et al. (2018), with slight modificaiton. The larvae were collected into a 15 ml centrifuge tube, and the excess fish water was removed. The larvae were washed twice with 1X PBS (without calcium and magnesium), and incubated in 1.5 ml of dissociation mix (1.44 ml 0.25% trypsin-EDTA + 60 μl Collagenase P at 50 mg/ml) was added. Mechanical dissociation was performed by vigorous pipetting with a P1000 tip for 30 seconds, followed by incubation at 30°C for 30 seconds. This step was repeated until the tissue was no longer visible. The dissociation was stopped by adding 2.4 ml Dulbecco’s Modified Eagle Medium (DMEM)-10% Fetal Bovine Serum (FBS) to the tube, followed by centrifugation at 500 *g* for 5 minutes at 4°C. The pellet should be visible at the bottom of the tube after centrifugation. After the pellet wash step, the cell suspension was filtered through a 40 μm cell strainer (#352350, Falcon, Corning, New York, USA) and proceeded with Reactive Oxygen Species (ROS) staining. For ROS detection, we used the CellROX Deep Red flow cytometry assay kit (#C10491, Invitrogen, Waltham, Massachusetts, USA), which is non-fluorescent in a reduced state but emits strong fluorescence upon oxidation. The staining was performed according to the user manual with slight modifications. The cells were incubated with 2.5 μM of CellROX solution in DMEM-10% FBS at 28.5°C for 30 minutes, protected from light. After staining, the cells were centrifuged, washed with 1X PBS, resuspended in sorting buffer (1X PBS-10% FBS), and filtered through a 40 μm cell strainer into a 5 ml tube, preparing them for flow cytometry analysis. Flow cytometry was performed using a BD FACSanato II (BD Biosciences, Franklin Lakes, New Jersey, USA). The gating strategies are followed the identification of viable cells, singlets, neutrophils, and CellROX signals in neutrophils. Histograms of the CellROX signal, with overlapping signals from different treatments, were generated using FCSalyzer Ver. 0.9.22. (https://sourceforge.net/projects/fcsalyzer/files/).

For neutrophil sorting, approximately 70 *Tg* (*mpx:EGFP*) larvae at 3 dpf per treatment were pooled and dissociated as described above. Neutrophils were isolated using a BD FACSAria™ III Cell Sorter (BD Biosciences) operating in purity mode, with gating strategies described as ROS analysis. Between 4,000 and 5,000 neutrophils were sorted directly into RNAzol® RT for subsequent RNA extraction.

### Whole-mount *in situ* hybridization (WISH)

WISH probes were generated by cloning 700-1000 bp of the target gene cDNA into the pGEM®-T Easy Vector (Promega). The plasmid was sequenced and linearized to synthesize antisense DIG-labeled RNA probes using the MEGAscript® SP6 or T7 Transcription Kit (Invitrogen, Waltham, MA). The resulting mixtures were purified with MicroSpin G-50 Columns (Cytiva, Buckinghamshire, UK) to remove unincorporated nucleotides. WISH was conducted following the method described by Thisse and Thisse (2008), with slight modifications. Proteinase K treatment conditions were as follows: 30 μg/ml for 15 minutes for 30 hpf embryos, 50 μg/ml for 25 minutes for 2 dpf embryos, and 70 μg/ml for 40 minutes for 3 dpf embryos. To minimize background noise during NBT/BCIP staining, embryos were incubated at 4°C. After staining, embryos were transferred to 100% methanol and stored overnight at −20°C to enhance staining contrast. Finally, embryos were mounted in glycerol for observation and image acquisition.

### Imaging and microscopy

Zebrafish larvae were examined using an inverted epifluorescence microscope (CKX41, Olympus, Tokyo, Japan) and photographed with a digital camera (EOS700D, Canon, Tokyo, Japan). Confocal microscopy images were captured using a Zeiss LSM 780 Confocal Microscope with 10X or 20X lenses. The imaging settings were standardized as follows: (1) Format: 512*512 or 1024*1024 pixels; (2) 488 nm laser for Alexa Fluor 488 and 561 nm laser for Alexa Fluor 568; (3) Pinhole size: 1AU, line average: 2; (4) Z-stack: 2.54 μm intervals with the 10X lens. Imaris Viewer (version: 10.2.0, RRID:SCR_007370) was used for 3-D reconstruction, Z-projection, and orthogonal view construction with maximum intensity of images.

### Statistics analysis

Statistical analyses were performed using Prism 8.0 (GraphPad Software). The data were subjected to parametric tests upon satisfying the assumptions of normality (Shapiro-Wilk test) and equality of variances (F-test). A two-tailed Student’s unpaired t-test was used for comparisons between the two groups. For comparisons involving three groups, a one-way ANOVA followed by Tukey’s post hoc test was performed.

Additionally, analyses involving both developmental stages and neutrophil counts across three groups utilized a two-way ANOVA with Dunnett’s multiple comparisons test.

## RESULTS

### Knockdown of Lpar1 causes neutrophil dispersal from hematopoietic tissue

To monitor neutrophil dynamics in live embryos, we employed the Tg*(mpx:EGFP)* transgenic line, which is widely used as a neutrophil-specific reporter and exhibits strong expression in mature neutrophils (Renshaw et al., 2006). We injected zebrafish embryos with 1.25 ng of standard MO (std MO) or Lpar1 splicing-blocking MO (Lpar1 MO) targeting the boundary between intron 2 and exon 2 (i2e2) of Lpar1 RNA that was alternatively spliced by skipping exon 2 and resulting in a truncated protein (Lee et al., 2008). In 30 hours post-fertilization (hpf), std MO-injected zebrafish embryos (will be called control embryos hereafter), neutrophils are primarily located in two hematopoietic tissues, the rostral blood island (RBI) and the caudal hematopoietic tissue (CHT) (Fig. 1A). Neutrophils were notably increased, well-organized and aligned with the CHT, with the remainder at the RBI and other regions in 2 days post-fertilization (dpf) embryos (dpf, Fig. 1B). In 3-dpf control larvae, neurophils remained at the CHT, with few scattered neurophils in the trunk region (Fig. 1C) . In contrast, Lpar1 MO-injected embryos (will be called Lpar1 morphants hereafter) appeared to have fewer neutrophils at 30 hpf (Fig. 1A). Strikingly, neurophils were all over the body, and few remained in the CHT at 2- and 3-dpf Lpar1 morphants (Fig. 1B, C). We counted the number of neutrophils in the dashed-enclosed trunk region of embryos (Fig. 1C, top image). The number of neutrophils in the trunk of Lpar1 morphants (19.8 ± 4.7) was significantly increased compared to that of control embryos (3.0 ± 2.2) (Fig. 1C). This scattered distribution of neutrophils and their reduced presence at the CHT suggest that neutrophils escape from the CHT in the loss of Lpar1.

**Figure 1.**
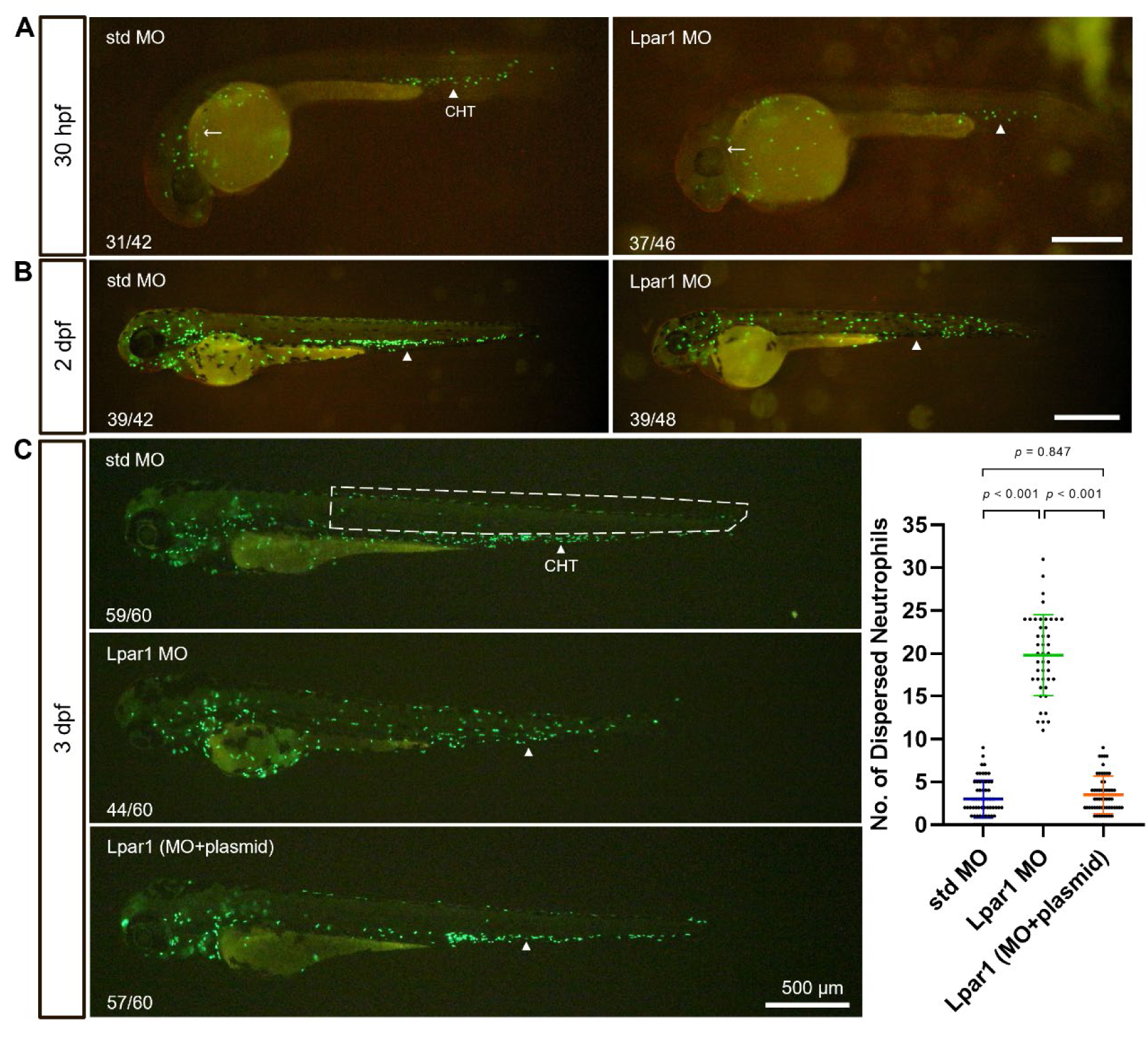
**Knockdown of Lpar1 causes the dispersal of neutrophils.** *Tg(mpx:EGFP)* zebrafish embryos were injected at the one-cell stage with 1.25 ng of either standard control MO (std MO) or Lpar1 splicing-blocking MO (Lpar1 MO). Embryos were cultured and photographed under bright and dark fields at designated times. Representative merged images are shown for embryos at 30 hours post-fertilization (hpf; A) and 2 days post-fertilization (dpf; B). The ratio of embryos exhibiting the depicted phenotype is indicated in the lower-left corner of each panel. EGFP-labeled neutrophils were observed in the caudal hematopoietic tissue (CHT, white arrowheads), yolk sac, rostral blood island (RBI, white arrows in A), and eye surroundings. Lpar1 MO injection led to the dispersion of neutrophils from the CHT. In a separate experiment, embryos were co-injected with Lpar1 MO and Lpar1 expression plasmid, examined, and shown as above. The number of neutrophils in the trunk region (outlined by dashed lines) was quantified at 3 dpf (C). Each group included 20 larvae across three independent experiments. Data are presented as mean ± s.d. and were analyzed using one-way ANOVA followed by Tukey’s post hoc test.

To verify whether the knockdown of Lpar1 specifically causes the dispersal of neutrophils, we co-injected the Lpar1 MO with a plasmid containing the coding region of Lpar1 in a pcDNA3.1/V5-His-TOPO expression vector (Lee et al., 2008). Ectopic expression of *lpar1* significantly decreased the number of dispersed neutrophils in Lpar1 morphants and restored the normal distribution pattern of neutrophils (Fig. 1C, bottom). These findings indicate that Lpar1 plays a crucial role in keeping neurophils in the CHT of zebrafish embryos.

### Lpar1 mutants also show neutrophil dispersal

To confirm the Lpar1 deficiency in neutrophil dispersal, we acquired a Lpar1 mutant line, sa38782, generated from the Zebrafish Mutation Project (Kettleborough et al., 2013). We obtained sa38782 embryos containing half/half *lpar1^+/+^* and *lpar1^+/sa38728^* genotype from the Zebrafish International Resource Center (ZIRC, https://zebrafish.org/home/guide.php). The sa38782 allele carries a C-to-T (C>T) mutation in Exon 2 of *lpar1*, which introduces a premature stop codon near the translation initiation site, leading to a predicted truncated Lpar1 protein consisting of only four amino acids (Fig. S1A). To identify the *lpar1^+/sa38728^*allele and validate the C>T mutation therein, we extracted the genomic DNA from the clipped tail fin of adults. We applied the derived Cleaved Amplified Polymorphic Sequence (dCAPS) method to design two forward primers introducing a cut site for a restriction enzyme (RE) AccI or ClaI in the PCR fragment of wild-type (WT) or sa38782 allele, respectively, and analyzed by capillary electrophoresis (Fig. S1B). The AccI can cut the WT strand to have two fragments of 209 and 25 base pairs (bp). The 25-bp peak was not detectable in capillary electrophoresis (Fig. S1C). In addition, the Sa38728 does not have the AccI cut site due to its C>T mutation. Thus, an intact 234-bp peak was found. On the other hand, a ClaI cut site was introduced in the Sa38728 strand that can cut the Sa38728 strand to two fragments of 211 and 24 bp (not detectable, Fig. S1C). An intact 235-bp peak was found in the WT strand with no ClaI cut site. The *lpar1^+/sa38728^*heterozygous zebrafish were identified by the presence of two peaks with 234 and 209 bp in the AccI-digested or 235 and 211 bp in the ClaI-digested PCR products (Fig. S1C). The above-described genotyping method was used throughout this study.

We crossed the identified homozygous *lpar1^sa23872/sa38728^*mutants with the *Tg (mpx:EGFP)* line to obtain the heterozygous and homozygous Sa23872 allele in the green fluorescent neutrophil background (will be called Lpar1 mutants hereafter unless otherwise stated). At 3 dpf, heterozygous Lpar1 mutants show a neutrophil distribution similar to that of WT embryos, but Lpar1 mutants exhibit high neutrophil dispersal similar to that observed in Lpar1 morphants (Fig. 2A). The number of dispersed neutrophils in the 3-dpf homozygous Lpar1 mutants (17.3 ± 6.2) are significantly higher than in heterozygous Lpar1 mutants (4.4 ± 3.2) and WT embryos (4.3 ± 2.7) (Fig. 2B). This increased neutrophil dispersal in homozygous Lpar1 mutants reinforces the importance of Lpar1 in the proper organization of neutrophil distribution in hematopoietic tissue.

**Figure 2.**
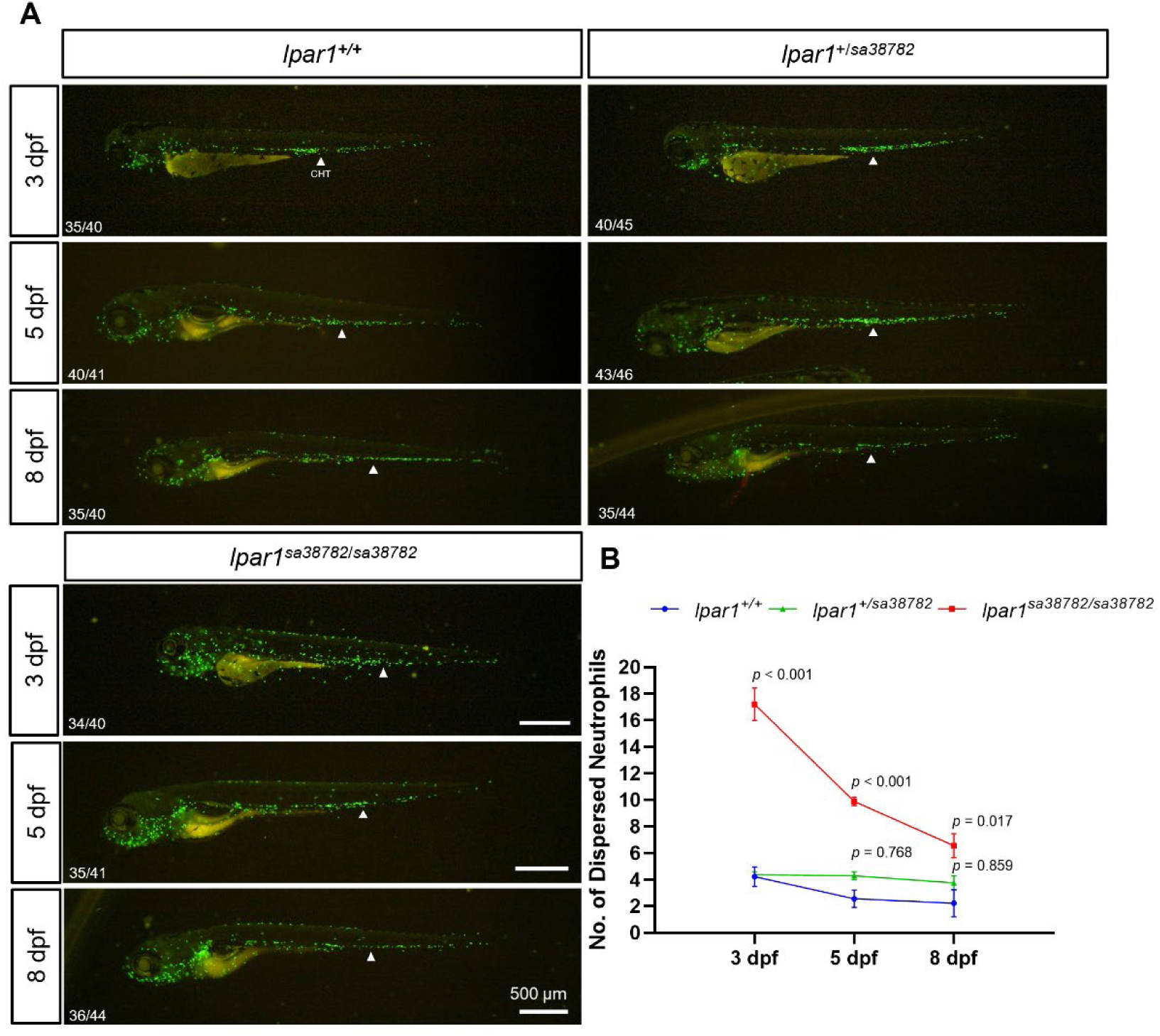
**Lpar1 sa38782 mutant phenocopies Lpar1 MO–induced neutrophil dispersal.** (A) Representative superimposed images of wild-type (WT, *lpar1_+/+_*) control, heterozygous mutant (*lpar1*_+/*sa38782*_), and homozygous mutant (*lpar1_sa38782_*_/*sa38782*_) larvae in Tg*(mpx:EGFP)* background at 3, 5, and 8 days post-fertilization (dpf) were captured under bright and dark fields. (B) The number of neutrophils dispersed from the caudal hematopoietic tissue (CHT, white arrowheads) was quantified as described in Fig. 1 in 10-16 larvae per group, across three independent experiments at different stages within WT, heterozygous mutant, and homozygous mutants. Values are shown as mean ± s.e.m. The Two-way ANOVA test with Dunnett’s multiple comparisons analyzed the distinction between the three genotypes across 3 to 8 dpf.

Lpar1 morphants develop pericardial and trunk edema at 5 dpf (Lee et al., 2008). We confirmed the development of edema in Lapr1 morphants, and also observed the recruitment of neutrophils to the edema sites at 5 dpf in the morphants but not in control embryos (Fig. S2). In contrast, no edema was observed in either heterozygous or homozygous Lapr1 mutants at 5 dpf or later stages. Interestingly, the number of dispersed neutrophils is still significantly higher in homozygous Lpar1 mutants than in other Lpar1 genotypes. However, the dispersed neutrophils in the trunk gradually reduce but still remain at a higher level in 5-dpf (Fig. 2A) and 8-dpf Lpar1 mutants (Fig. 2 B). This finding provides strong genetic evidence for a role of Lpar1 in the distribution of neutrophils, and this regulation is independent of edema.

### Knockdown of Lpar1 causes macrophage dispersal from hematopoietic tissue

Macrophages are an essential component of the innate immune system that can resolve inflammation and apoptotic neutrophils or assist in the reverse migration of neutrophils (Silva, 2010; Tauzin et al., 2014). We therefore asked whether macrophages also exhibit a dispersed distribution in response to neutrophil dispersal. To investigate the role of Lpar1 in macrophage distribution, we employed the *Tg(mpeg1:EGFP)* reporter line to examine macrophage dynamics in live embryos.

In the 3-dpf control larvae, most macrophages were localized within the CHT, while few peripheral macrophages were in the trunk region, denoted as tissue-resident macrophages. Conversely, an increase in macrophages was observed in the trunk region of Lpar1 morphants, accompanied by a corresponding decrease in the CHT (Fig. S3).

Lpar1 morphants show a significantly higher number of macrophages (9.3 ± 4.3) in the trunk, including tissue-resident macrophages, compared to that in control embryos (6.0 ± 4.3) (p = 0.019; Fig. S3). These findings suggest that Lpar1 also influences macrophage dynamics, although the extent of dispersal appears less evident compared to neutrophils.

### Elevated epidermal apoptosis drives neutrophil dispersal in Lpar1 morphants

We observed dispersed neutrophils appear near the surface of the fish under epi-fluorescence microscopy. To determine the precise location of dispersed neutrophils in Lpar1 morphants. We crossed the *Tg* (*mpx: EGFP*) zebrafish and *Tg* (*krt4*: *h2afv-mCherry*)^cy9^ zebrafish expressing mCherry (Chen et al., 2014). The resulting double-transgenic embryos allow us to visualize the neutrophils in green fluorescence and superficial epidermal cells (SECs) in red fluorescence simultaneously. Single optical sections captured by confocal microscopy revealed that dispersed neutrophils in Lpar1 morphants were frequently in close contact with SECs, whereas most neutrophils in the control embryos remained within the CHT (Fig. 3A). To quantify this observation, we measured the percentage of dispersed neutrophils in contact with SECs relative to the total number of dispersed neutrophils, and found that 84.8 ± 2.4% of dispersed neutrophils in Lpar1 morphants were localized near SECs that was significantly higher than that (14.7 ± 7.7%) in control embryos (Fig. 3B). These findings suggested that Lpar1 deficiency promotes infiltration and recruitment of neutrophils into the superficial epidermal layer during early development.

**Figure 3.**
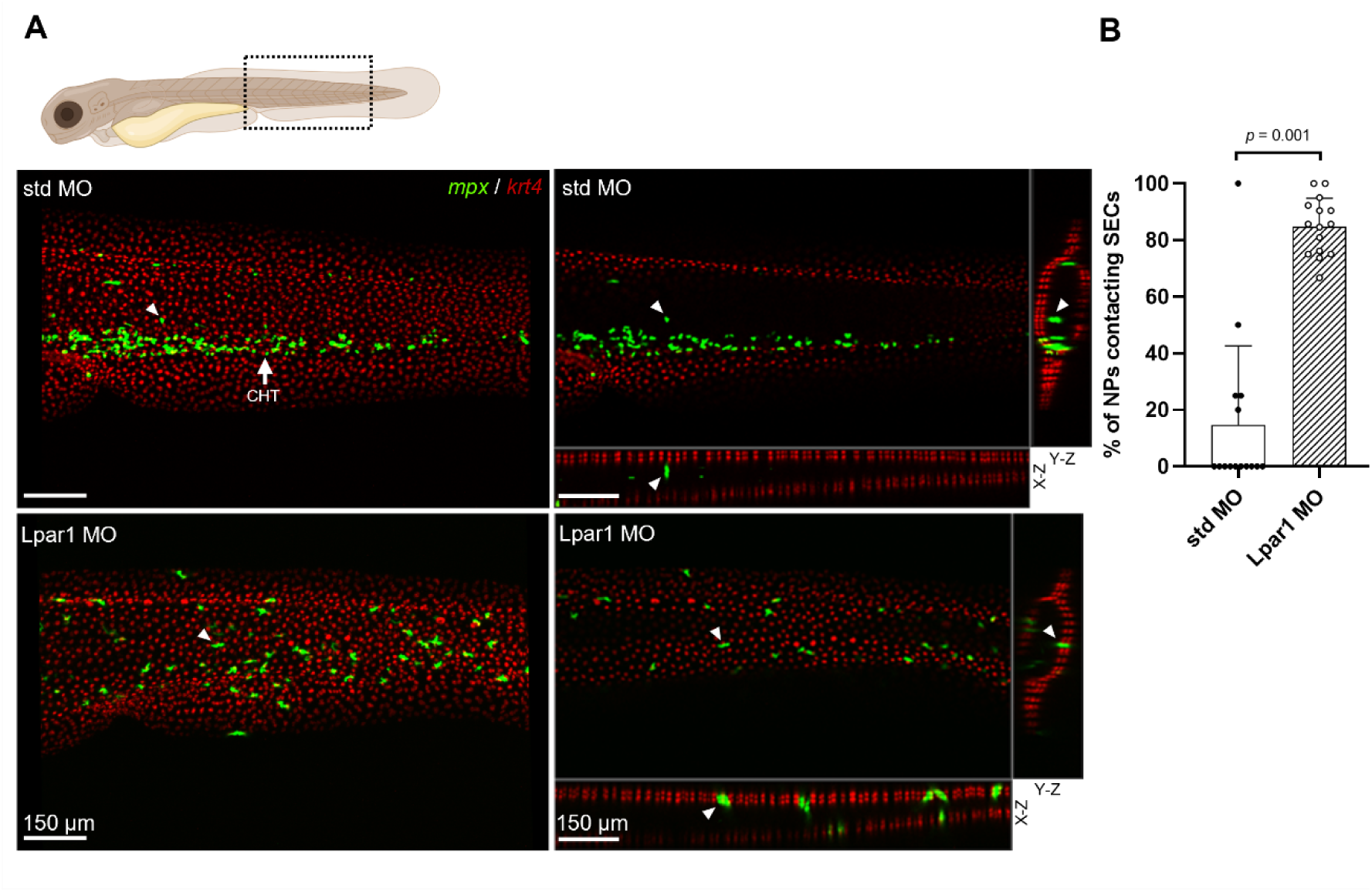
**Dispersed neutrophils in Lpar1 morphants are in close contact with the superficial epidermal cells.** Double transgenic zebrafish embryos from the cross of Tg*(mpx:EGFP)* and Tg*(krt4:h2afv-mCherry)^cy9^* were microinjected at the one-cell stage with 1.25 ng of either standard control (std) or Lpar1 splicing-blocking morpholino oligonucleotides (MO). Injected embryos were cultured to 3 days post-fertilization (dpf), subjected to whole-mount immunohistochemistry using antibodies against EGFP and mCherry, and imaged under confocal microscopy. (A) Representative lateral views of maximum projection at the box region are shown in the cartoon (top). Confocal microscopy images show neutrophils (green and indicated by white arrows) dispersed from the caudal hematopoietic tissue (CHT) in both control and morphant embryos. Corresponding single optical sections and orthogonal views from the same images, with white arrows indicating the same neutrophils at lateral views, demonstrate that dispersed neutrophils in morphants but not std MO-injected embryos are in close contact with superficial epidermal cells (SECs, krt4^+^ cells in red). (B) Quantification of dispersed neutrophils in contact with SECs was performed on 5 larvae per group, across 3 independent experiments, and presented as percentages. Data are shown as mean ± s.d., and analyzed using Student’s unpaired t-test.

To investigate the mechanisms underlying neutrophil dispersal, we utilized Lpar1 morphants for further analysis, which exhibited an earlier onset and a more severe phenotype compared to the mutants. The LPA-Lpar1 signaling pathway has been shown to mediate cell proliferation, differentiation, and keratinocyte barrier function in murine or human cell culture models (Kim et al., 2021; Sumitomo et al., 2019). Since most dispersed neutrophils in the 3-dpf Lpar1 morphants were in close contact with SECs, we hypothesized that epidermal homeostasis is affected without Lpar1. Thus, we performed a TUNEL assay in Tg(*krt4: h2afv-mCherry*)^cy9^ embryos to visualize cell apoptosis in the epidermal layer. We examined apoptosis in Lpar1 morphants at 33 hpf, before the occurrence of neutrophil dispersal. A few apoptotic SECs in the trunk region were noted in the control embryos, which were presumed to be programmed cell death during embryogenesis (Fig. 4A, A’). On the contrary, significantly more apoptotic SECs were found in the trunk of Lpar1 morphants, suggesting that Lpar1 is involved in SEC survival during early development (31.7 ± 11.2 in Lpar1 morphants versus 15.3 ± 8.0 in the control embryos; *p* = 0.023; Fig. 4A-A’’). These apoptotic cells could release damage-associated molecular patterns (DAMPs), recruiting neutrophils or macrophages to clear cell debris and help remodel tissue homeostasis (Pittman & Kubes, 2013).

**Figure 4.**
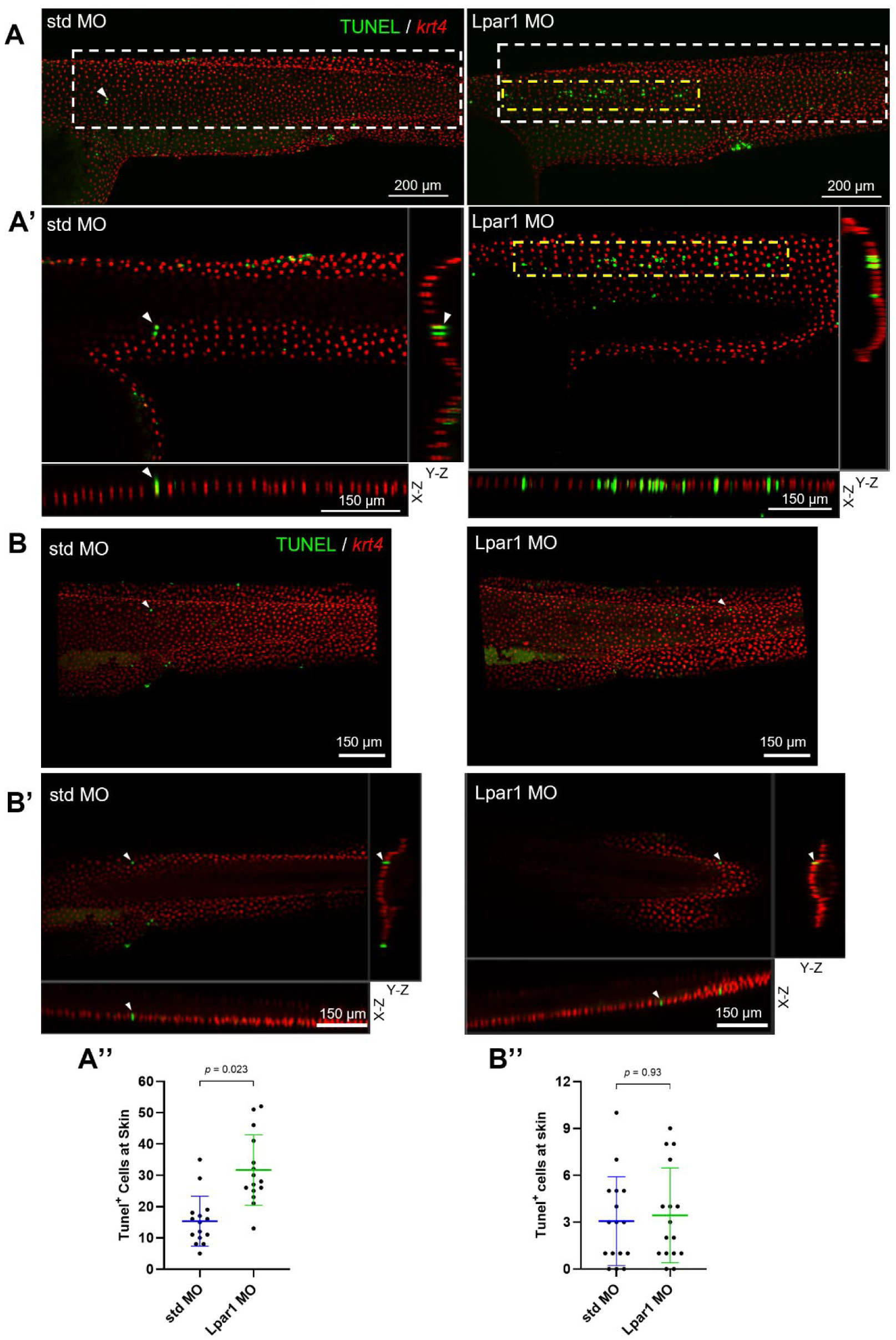
Knockdown of Lpar1 promotes apoptosis in superficial epidermal cells at 33 hpf. Tg(*krt4:h2afv-mCherry*)_cy9_ zebrafish embryos were injected at the one-cell stage with 1.25 ng of either standard control MO (std MO) or Lpar1 MO (Lpar1 MO). Embryos were cultured to either 33 hours post-fertilization (hpf; A, A’, A’’) or 3 days post-fertilization (dpf; B, B’, B’’), followed by TUNEL assay and whole-mount immunohistochemistry using anti-mCherry antibodies. (A) Representative lateral maximum intensity projections show increased green TUNEL_+_ cells in Lpar1 MO-injected embryos (right, yellow dashed boxes) compared to std MO-injected controls (left, white arrowheads). (A’) Single optical sections with corresponding X-Z and Y-Z views highlight TUNEL_+_ signals within the superficial epithelial cell (SEC) layer. (B, B’) At 3 dpf, the number of TUNEL_+_ cells appeared reduced in both std MO-and Lpar1 MO-injected embryos. (A’’, B’’) Quantification of TUNEL_+_ cells within the white boxed regions was performed on 5 larvae per group from 3 independent experiments. Data are presented as mean ± s.d. and analyzed using Student’s unpaired *t*-test.

Therefore, we examined apoptosis in the larvae at 3 dpf, when neutrophil dispersal was at the highest level. There were only sparse apoptotic cells in control larvae and Lpar1 morphants, suggesting effective cleanup of apoptotic cells (Fig. 4B-B’’).

To further assess whether excessive epidermal apoptosis triggers neutrophil dispersal in morphants, we treated embryos with the pan-caspase inhibitor zVAD-fmk to block apoptosis. Embryos were incubated with 100 μM zVAD-fmk from 32 hpf to 3 dpf. Although zVAD-fmk treatment slightly reduced the number of randomly dispersed neutrophils in control embryos, it partially rescued the neutrophil dispersal phenotype in Lpar1 morphants (12.5 ± 6.6 in morphants treated with inhibitor versus 24.0 ± 9.8 in morphants; *p* = 0.0004; Fig. 5D). These results suggest that elevated epidermal apoptosis is a key contributor to neutrophil dispersal in morphants.

**Figure 5.**
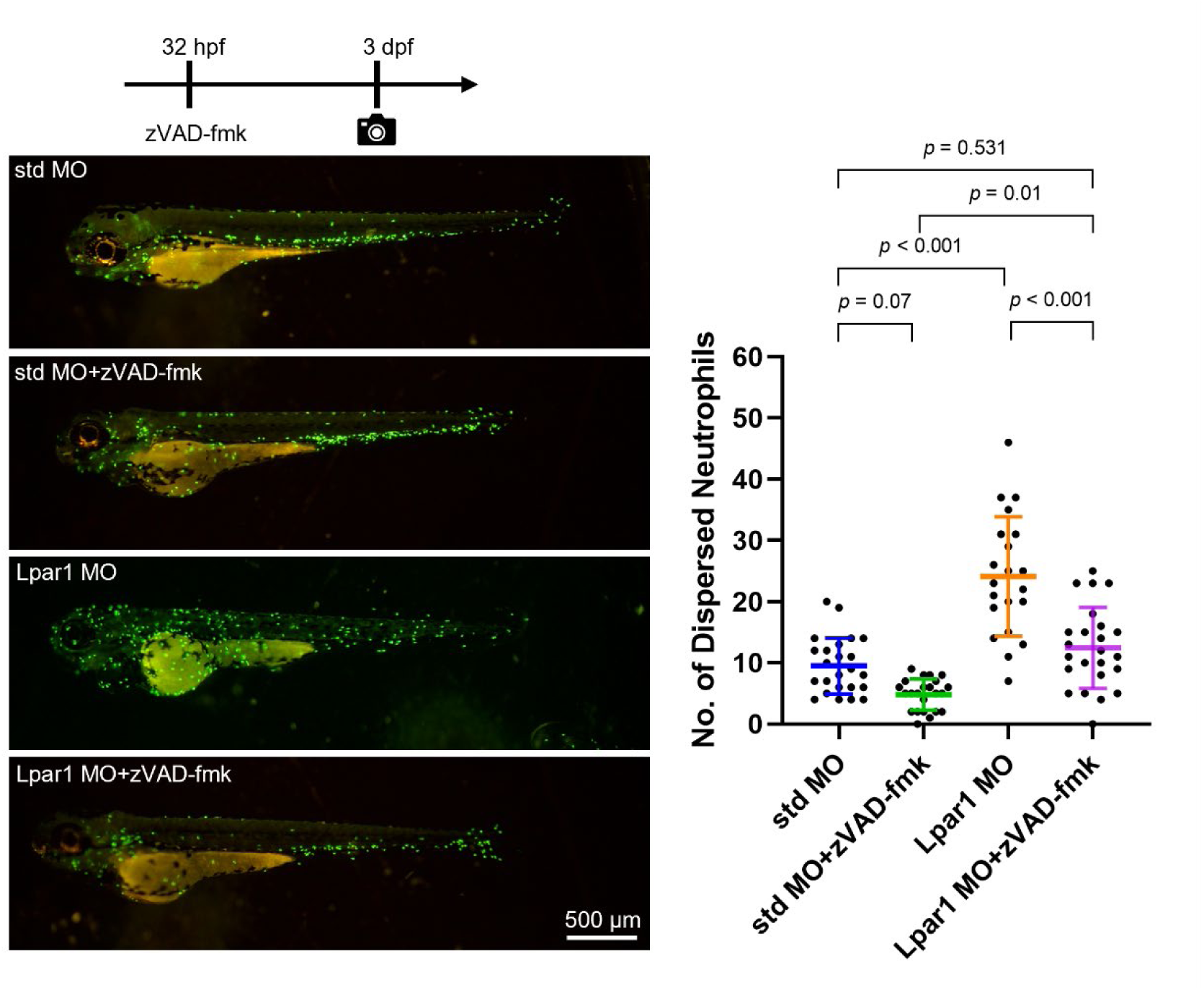
**Inhibition of apoptosis partially rescues neutrophil dispersal in Lpar1 morphants.** Embryos were injected at the one-cell stage with 1.25 ng of either standard control MO (std MO) or Lpar1 MO (Lpar1 MO), treated with 100 μM ZVAD-fmk at 32 hours post-fertilization (hpf), and imaged at 3 days post-fertilization (dpf). Dispersed neutrophils were quantified at 3 dpf in 7–9 larvae per group across three independent experiments. Data are presented as mean ± s.d. and analyzed using one-way ANOVA followed by Tukey’s post hoc test.

### Elevated expression of *tnfr2* in Lpar1 morphants and *il1b* in neurophils

Since overall cell apoptosis in Lpar1 morphants was similar to that in control larvae at 3 dpf, neutrophil dispersal should also have resolved by this stage. However, neutrophil dispersal was at its most severe level at 3 dpf, suggesting that other factors may prevent dispersed neutrophils from reverse migrating to the CHT or undergoing apoptosis.

Unresolved tissue inflammation may create a microenvironment that retains neutrophils. Therefore, we examined the mRNA expression of inflammatory genes—*cxcl8a*, *il1b*, *tnfr2*, and *duox1* in the whole control and Lpar1 morphant larvae at 3 dpf by RT-qPCR analysis. Cxcl8a is a small cytokine upregulated in wound tissue and acts as an important chemoattractant molecule guiding neutrophil recruitment (de Oliveira et al., 2013). Interleukin 1 beta (*il1b*) and tumor necrosis factor receptor superfamily member 1B (*tnfrsf1b*, also known as *tnfr2*) are pro-inflammatory genes known to be upregulated in inflamed tissue, and TNF-TNFR2 signaling is further involved in neutrophil activation during bacterial skin infection (Xie et al., 2020; Youn et al., 2023). Dual oxidase (*duox1*) is an NADPH oxidase that generates hydrogen peroxide (H_2_O_2_), recruiting neutrophils that detect the H_2_O_2_ gradient through Lyn, a redox sensor on neutrophils (Niethammer et al., 2009; Yoo et al., 2011). We found that only *tnfr2* was significantly upregulated in Lpar1 morphants (Fig. 6A). While *cxcl8a* and *il1b* were slightly upregulated, and *duox1* was slightly downregulated, they were not significant compared to the control group (Fig. 6A). Although apoptotic SECs were removed in 3-dpf Lpar1 morphants, the inflammatory *tnfr2* remained upregulated, indicating that the inflammation had not yet resolved.

**Figure 6.**
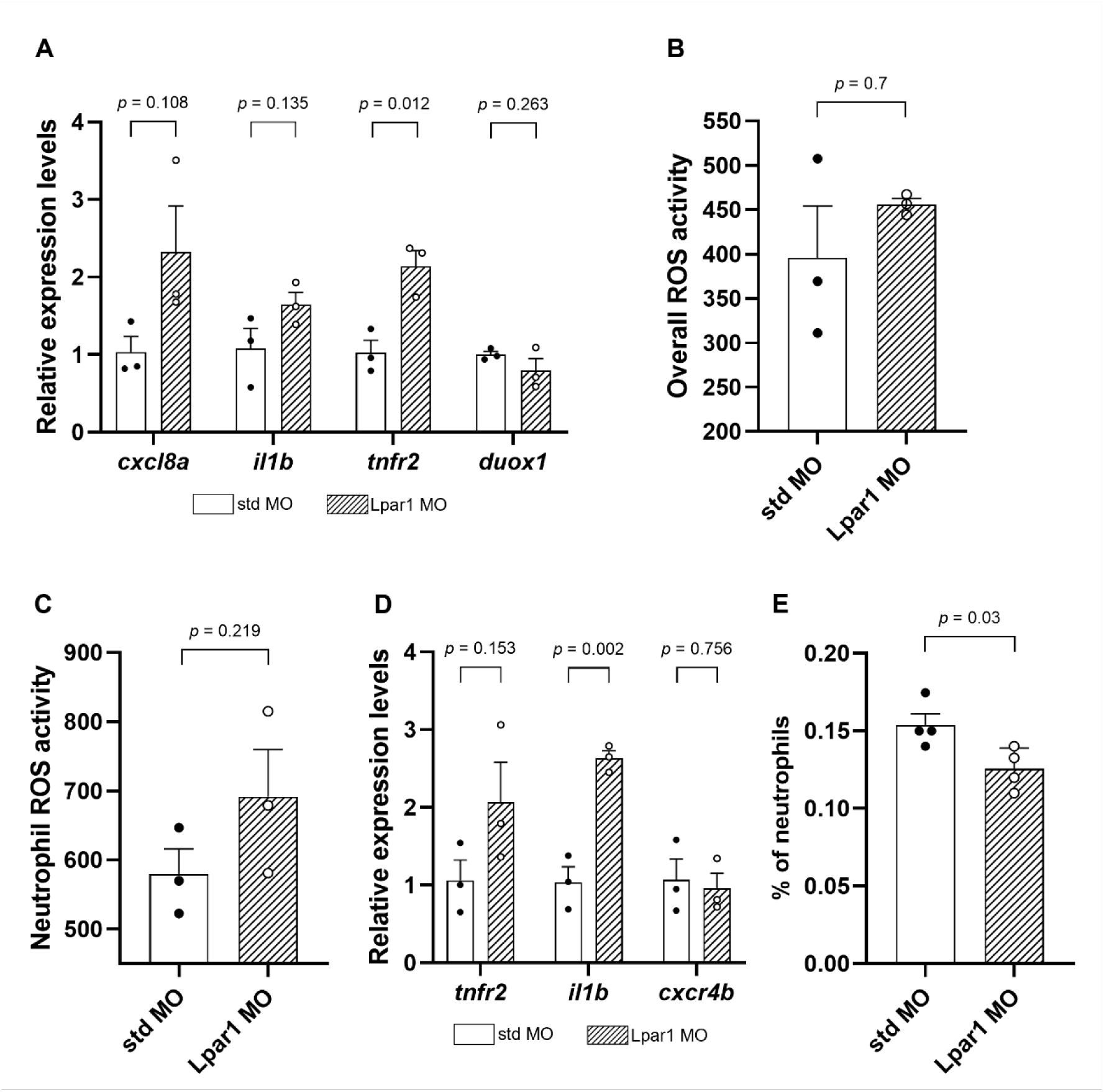
**Knockdown of Lpar1 elevates pro-inflammatory gene expression and reduces neutrophil abundance without significantly altering ROS activity in zebrafish larvae** (A) Relative mRNA expression of *cxcl8a*, *il1b*, *tnfr2*, and *duox1* in 3-dpf larvae following morpholino knockdown. Larvae were injected at the one-cell stage with 1.25 ng of either standard control morpholino oligonucleotides (std MO) or Lpar1 splicing-blocking morpholino oligonucleotides (Lpar1 MO). Expression was quantified by RT-qPCR using RNA pooled from 40 larvae per condition for each biological replicate. Values were normalized to the reference gene, *eef1a1l1,* and the fold change level was calculated using the 2^−ΔΔCT^ method. Data are presented as mean ± s.e.m. from 3 independent biological replicates. The unpaired Student’s t-test revealed a significant difference in *tnfr2* expression level between control and Lpar1 morphants (*p* = 0.012). (B-E) Tg(*mpx:EGFP*) embryos were injected with std MO or Lpar1 MO at the one-cell stage and raised to 3 dpf. Larvae were then dissociated into single-cell suspensions and subjected to flow cytometry for neutrophil sorting or analysis. (B, C) Reactive Oxygen Species (ROS) activity was assessed using CellROX Deep Red staining in single-cell suspensions at 3 dpf. (B) Whole-larva ROS activity was quantified in singlet populations based on fluorescence intensity and shown as mean intensity and s.e.m. (C) Neutrophil ROS activity is determined by gating on the EGFP-positive population and is shown as mean and s.e.m. Unpaired Student’s t-tests showed no significant difference (*p* > 0.05) in ROS activity between the control group and Lpar1 morphants for either whole larvae (B) or neutrophils (C). (D) Relative expression of *tnfr2*, *il1b*, and *cxcr4b* in sorted neutrophils (EGFP-positive cells) at 3 dpf. RNA was extracted from pooled neutrophils per condition for each biological replicate. RT-qPCR data are presented as mean ± s.e.m. from three independent experiments. A significant upregulation of *il1b* was detected in neutrophils from Lpar1 morphants (*p* = 0.002). (E) The proportion of neutrophils (GFP-positive cells; NPs) relative to total analyzed cells was quantified by flow cytometry. Data are presented as mean ± s.e.m. from four independent biological replicates. An unpaired Student’s t-test revealed a significant reduction in the neutrophil population in Lpar1 morphants compared to controls (*p* = 0.03).

Since the upregulated *tnfr2* indicated the unresolved inflammation in Lpar1 morphants at 3 dpf, we wondered whether neutrophils were activated in response to the inflammation. Activated neutrophils rapidly generate and release reactive oxygen species (ROS) in response to inflammatory mediators or pathogen components, a process known as the oxidative burst or respiratory burst (Lambeth, 2004). The examination of oxidative burst in neutrophils is commonly used to identify activated neutrophils (Chen & Junger, 2012). To assess ROS activity, we performed a ROS assay using CellROX, a general probe that detects ROS and emits fluorescence upon oxidation, in Tg(*mpx:EGFP*) larvae at 3 dpf. ROS activity in neutrophils was analyzed by flow cytometry, and the neutrophil population was defined based on EGFP fluorescence intensity and cell size (Fig. S4A). First, the overall ROS activity in Lpar1 morphants was slightly increased compared to controls, but the difference was not statistically significant (Fig. 6B; Fig. S4B). The similar overall ROS levels were consistent with the unchanged expression of *duox1*. In addition, ROS activity within neutrophils was also slightly elevated in Lpar1 morphants but not significantly different from that in the control group (Fig. 6C; Fig. S4C).

Since ROS levels remained similar in both groups, we next asked whether pro-inflammatory gene expression specifically in neutrophils was altered in Lpar1 morphants. To determine whether the absence of Lpar1 might affect neutrophil physiology, we first examined whether Lpar1 is expressed in neutrophils. Neutrophils were isolated from 3 dpf Tg(*mpx:EGFP*) larvae by flow cytometry, sorting for EGFP-positive cells. Indeed, we detected *lpar1* expression in the sorted neutrophils (Fig. S5), suggesting it may play a direct role in neutrophil regulation. We then examined the expression of pro-inflammatory markers *il1b* and *tnfr2* in neutrophils. The *il1b* was significantly upregulated in neutrophils of Lpar1 morphants (Fig. 6D), whereas *tnfr2* showed a modest but non-significant increase compared to controls. Last, we evaluated the expression of *cxcr4b*, the receptor for Cxcl12a, which regulates neutrophil retention in hematopoietic tissue. Its expression was unchanged between groups (Fig. 6D).

We further analyzed the ratio of neutrophils to total cells based on flow cytometry data, providing an estimate of overall neutrophil numbers in the larvae. The results showed that Lpar1 knockdown led to a significant reduction in total neutrophil abundance at 3 dpf (Fig. 6E). This finding is consistent with the distribution pattern observed in fluorescent imaging (Fig. 1C) and suggests that myelopoiesis may also be impaired in Lpar1 morphants.

### Neutrophil dispersal in Lpar1-deficient embryos resulting from downregulation of *cxcl12a* retention signal in hematopoietic tissue

Regarding what drives neutrophils to disperse from CHT, in addition to the plausible inflammatory signals from the epidermis, other factors might promote the release of neutrophils from the CHT. The CXCL12/CXCR4 axis, crucial for neutrophil retention in mammalian bone marrow, is also essential for retaining neutrophils within the CHT of zebrafish. Impairment of this signaling in zebrafish can lead to the unregulated release of neutrophils into the blood circulation (Paredes-Zuniga et al., 2017; Walters et al., 2010). Cxcl12a (also known as Sdf1a) is highly expressed in regions where neutrophils are formed, including the CHT in zebrafish. Therefore, we hypothesized that the knockdown of Lpar1 would reduce the expression level of *cxcl12a* in the CHT.

We examined the mRNA expression pattern of *cxcl12a* by whole-mount *in situ* hybridization (WISH) from 30 hpf to 3 dpf. *cxcl12a* was mainly expressed in the head, midline, pronephric duct, and CHT at 30 and 48 hpf (Fig. 7A, B, left), as described in previous research (Torregroza et al., 2012; Walters et al., 2010). In Lpar1 morphants, we found a sharply reduced expression of *cxcl12a* in the pronephric duct region and CHT at both 30 and 48 hpf (Fig. 7A, B, right). By 3 dpf, the expression pattern of *cxcl12a* had shifted, with expression in the dorsal aspect of the posterior cardinal vein (PCV) (Fig. 7C, left) as previously reported (Cha et al., 2012). In addition, *cxcl12a* is also expressed in midline tissue and CHT at this stage. In Lpar1 morphants, the expression of *cxcl12a* in PCV and CHT showed an indistinguishable signal compared to the control group (Fig. 7C, right). Moreover, we noted that *cxcl12a* was strongly expressed in the midline in Lpar1 morphants compared to moderate expression in control embryos from 2 dpf to 3 dpf (Fig. 7 B, C). Taken together, the retention signal of neutrophils at the hematopoietic tissue, *cxcl12a*, was downregulated in the absence of Lpar1. This reduced expression occurred before the dispersal of neutrophils, suggesting that the loss of the retention signal at the CHT might also be a causative factor for neutrophil dispersal.

**Figure 7.**
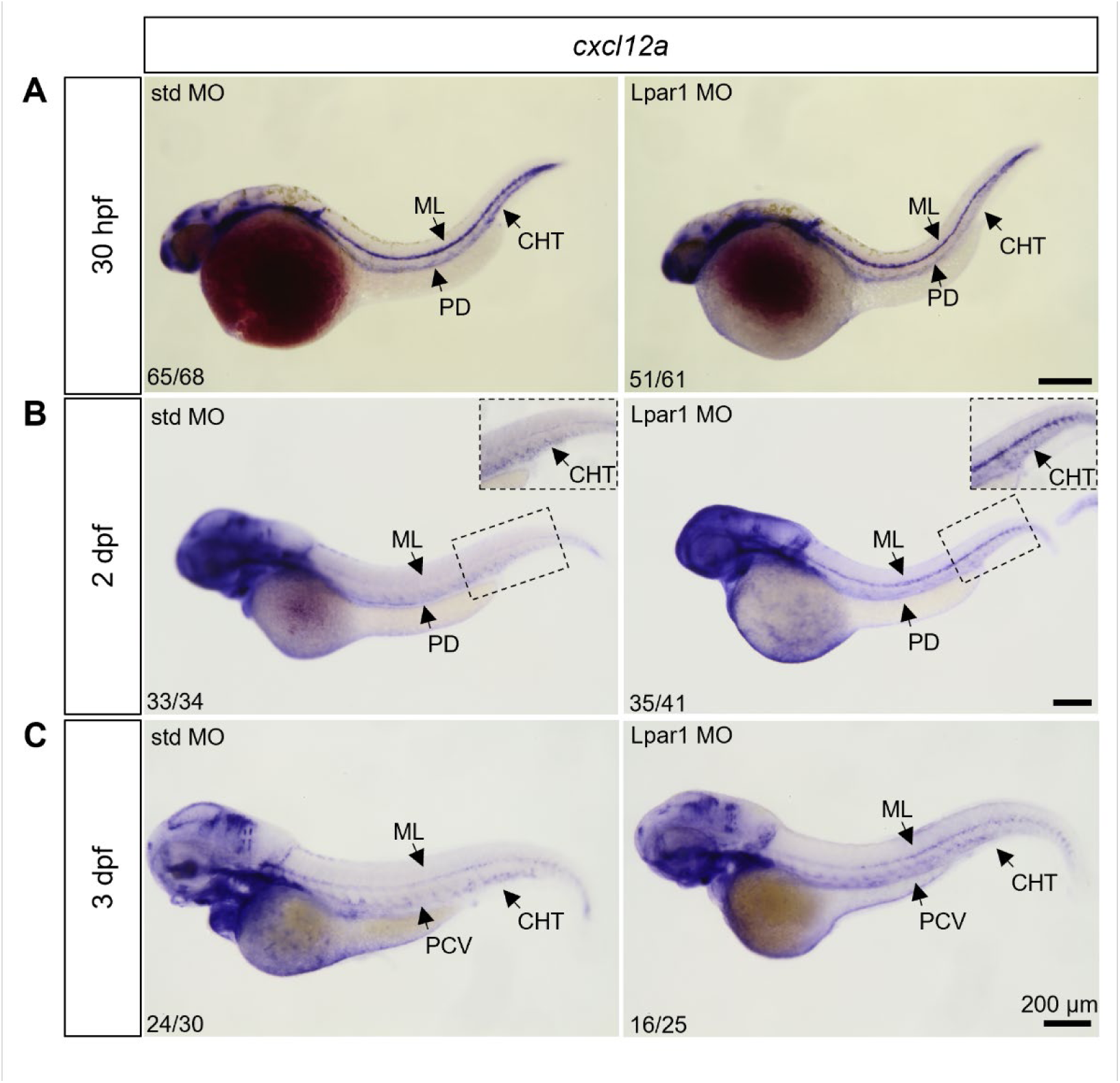
**Knockdown of Lpar1 downregulates the expression of cxcl12a in the hematopoietic tissue of zebrafish.** Embryos were injected at the one-cell stage with 1.25 ng of either standard control MO (std MO) or Lpar1 splicing-blocking MO (Lpar1 MO), fixed at (A) 30 hours, (B) 2 days, (C) 3 days post-fertilization (30 hpf, 2 and 3 dpf), and subjected to whole-mount *in situ* hybridization (WISH) against *cxcl12a*. One representative image is shown for each treatment at designated times. CHT: caudal hematopoietic tissue; ML: midline tissue; PD: pronephric duct; PCV: posterior cardinal vein.

To functionally assess whether reduced *cxcl12a* expression in the CHT contributes to neutrophil dispersal, we ectopically expressed *cxcl12a* by injecting mRNA at the one-cell stage. We first tested whether restoring *cxcl12a* levels could suppress neutrophil dispersal in Lpar1 morphants. Coinjection of 200 pg *cxcl12a* mRNA with Lpar1 MO significantly reduced neutrophil dispersal (18.6 ± 6.7) compared to morphants alone (27.9 ± 9.7) (*p* = 0.0002; Fig. S6), indicating a partial rescue. We next asked whether the same rescue could be achieved in homozygous Lpar1 mutants. Surprisingly, a lower dose of *cxcl12a* mRNA (100 pg) was sufficient to significantly reduce neutrophil dispersal in mutants (12.2 ± 6.1 in *cxcl12a* ectopically expressed mutants versus 22.4 ± 8.8 in mutants; *p* = 0.0005; Fig. 8). This suggests that the mutant background may be more sensitive to restored *cxcl12a* levels, possibly due to a milder dispersal phenotype. Additionally, in both Lpar1 morphants and mutants, ectopic *cxcl12a* expression shifted dispersed neutrophils toward the dorsal trunk region, contrasting with the scattered distribution seen in untreated groups. Together, these findings demonstrate that Cxcl12a functions as a key retention signal in the CHT, and that restoring its expression can partially reverse the dispersal phenotype in Lpar1-deficient embryos.

**Figure 8.**
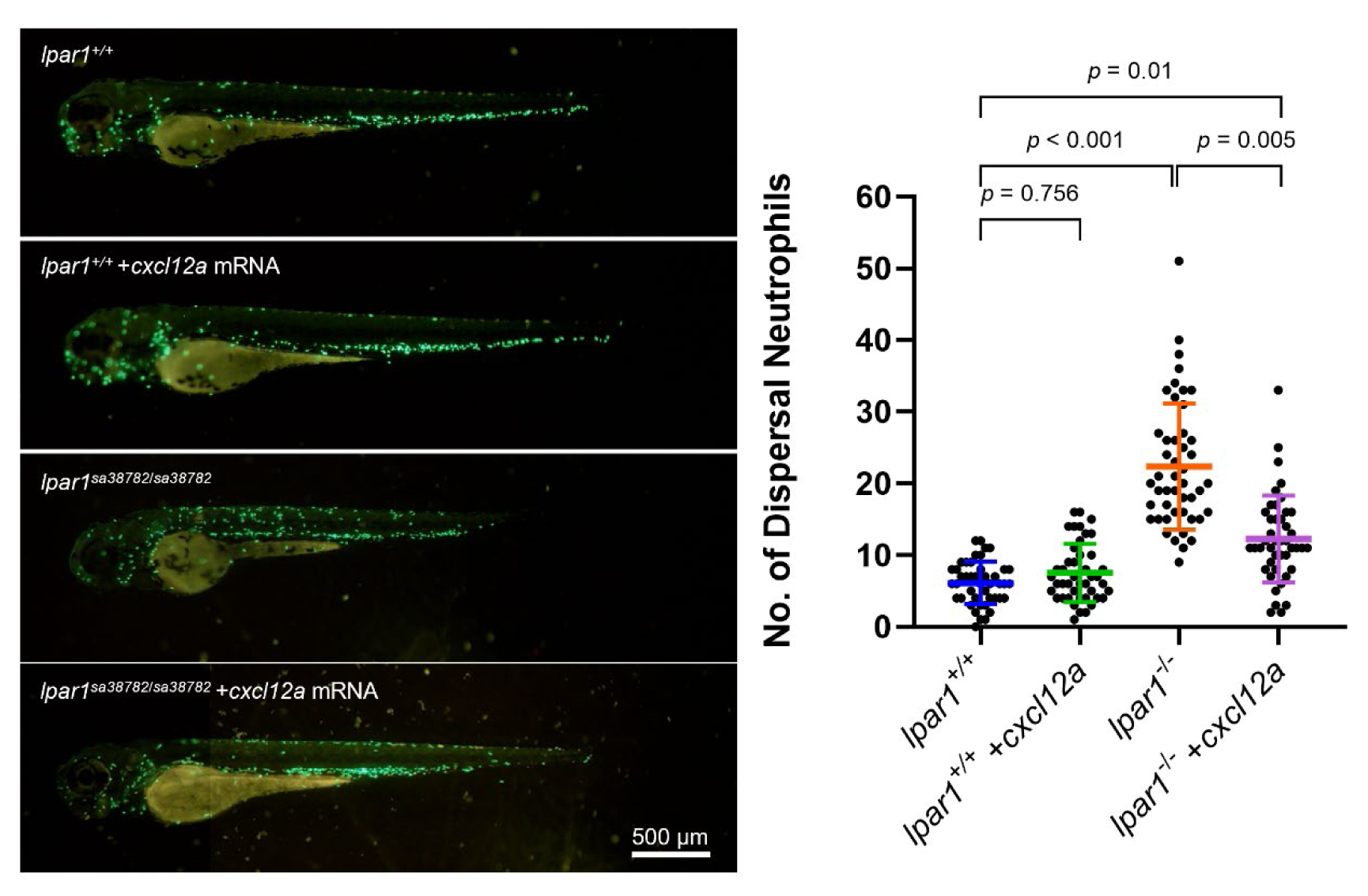
**Ectopic expression of cxcl12a ameliorate the neutrophil dispersal in Lpar1 mutants.** Embryos were injected at the one-cell stage with 100 pg of *cxcl12a* mRNA into either wild-type or homozygous mutant backgrounds and cultured until 3 days post-fertilization (dpf). Representative composite images of wild-type (*lpar1*_+/+_) controls and homozygous mutants (*lpar1*_sa38782/sa38782_) in the Tg(*mpx:EGFP*) background were captured at 3 dpf under both bright-field and fluorescence microscopy. Images were compiled from different focal planes to highlight the head and trunk regions. Dispersed neutrophils were quantified at 3 dpf using 15–16 larvae per group from three independent experiments. Data are presented as mean ± s.d. and analyzed using one-way ANOVA followed by Tukey’s post hoc test.

## DISCUSSION

### LPA-LPAR1 signalling is essential for the retention of neutrophils in the caudal hematopoietic tissue

LPA–LPAR1 signaling has been implicated in mediating leukocyte infiltration in various chronic inflammatory conditions, and disruption of this pathway has been shown to alleviate excessive neutrophil recruitment. However, its role in regulating neutrophil behavior under steady-state conditions or during early vertebrate development remains unexplored. In this study, we investigated the function of Lpar1 in neutrophil dynamics using both MO knockdown and a loss-of-function zebrafish mutant line. We found that loss of Lpar1 led to pronounced dispersal of neutrophils and macrophages from the CHT, a key site for myeloid retention during early development. Prior to the onset of neutrophil dispersal, Lpar1-deficient embryos exhibited increased apoptosis in superficial epidermal cells and reduced *cxcl12a* expression within the CHT. These early defects likely compromise tissue homeostasis and the retention environment, facilitating neutrophil infiltration into the epidermis and promoting local inflammation (Fig. 9).

**Figure 9.**
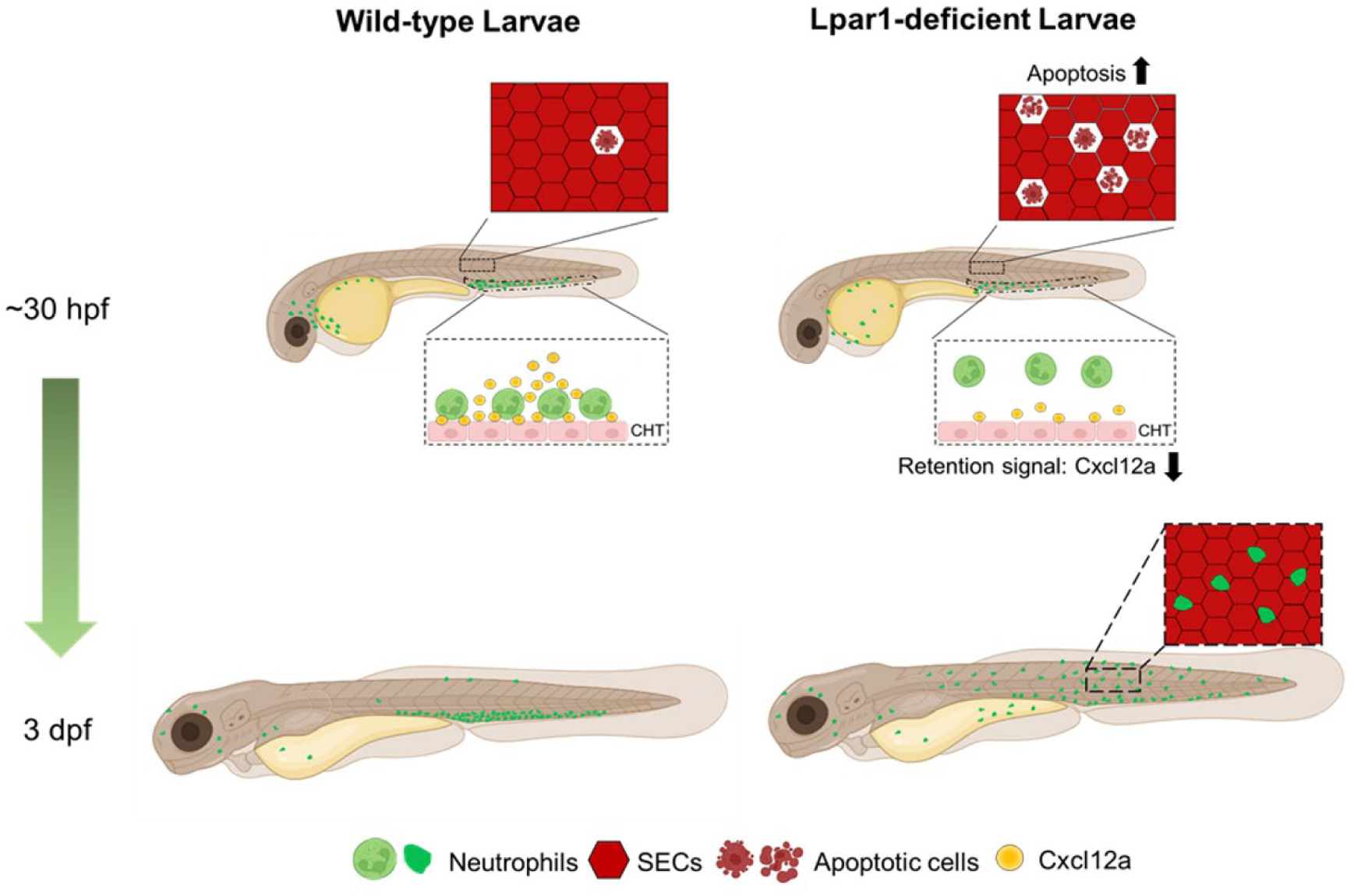
**Proposed model for neutrophil dispersal from caudal hematopoietic tissue (CHT) in Lpar1-deficient zebrafish.** Loss of Lpar1 function leads to increased apoptosis of superficial epidermal cells (SECs), which in turn triggers local inflammation that promotes neutrophil infiltration. Concurrently, *cxcl12a* expression is reduced in the CHT, diminishing the retention signals for neutrophils. As a result, neutrophils escape from the CHT and migrate toward the inflamed skin. By 3 days post-fertilization (dpf), Lpar1-deficient larvae show significantly enhanced neutrophil extravasation from the CHT and accumulation in the epidermis.

### Temporal regulation of neutrophil dynamics by Lpar1 during development

While we observed significant neutrophil dispersal in both Lpar1 morphants and mutants at 3 dpf, the dispersal pattern differed slightly between the two models. Mutant embryos had more neutrophils within the CHT than morphants, despite only a slight difference in the number of dispersed neutrophils. Given our data showing a reduced total neutrophil count in morphants, this distribution discrepancy may reflect impaired myelopoiesis in Lpar1 morphants. These findings suggest that mutants might better tolerate Lpar1 deficiency in neutrophil production compared to morphants. However, because our reporter line is driven by the *mpx* promoter, which is highly expressed in intermediate progenitors and mature neutrophils (Jin et al., 2012; Kirchberger et al., 2024). Therefore, we cannot exclude the possibility that Lpar1 deficiency, particularly in morphants, impairs neutrophil maturation, leading to developmental arrest at a progenitor stage that is minimally detected in this reporter line.

Notably, neutrophil dispersal gradually resolved by 8 dpf in Lpar1 mutants but not in morphants. This could be due to the onset of pericardial and trunk edema in morphants at 5 dpf, which was absent in mutants. We previously reported that this edema results from defective thoracic duct formation, disrupting body fluid homeostasis (Lee et al., 2008). Assessing whether the thoracic duct forms properly in Lpar1 mutants could help explain the differential edema phenotypes. The resolution of neutrophil dispersal in mutants may also relate to the developmental shift of definitive myelopoiesis from the CHT to the kidney. In zebrafish, renal tubules express high levels of Cxcl12a, which serves as a homing cue for hematopoietic stem cells (Glass et al., 2011). Therefore, it is plausible that homing signals released by the developing kidney contribute to the resolution of neutrophil dispersal in Lpar1 mutants. We observed a gradual accumulation of neutrophils in the pronephros (the embryonic precursor to the kidney) between 5 and 8 dpf, suggesting that chemotactic signals from this region remain functional. This supports the idea that Lpar1 is primarily required during the early stages of myelopoiesis.

### Apoptotic epidermal cells as triggers of neutrophil dispersal and potential links to psoriasis-like pathology

To investigate potential triggers of neutrophil dispersal, we observed a significantly higher number of apoptotic superficial epidermal cells (SECs) prior to dispersal in Lpar1 morphants. Additionally, most dispersed neutrophils were in close contact with SECs at 3 dpf in morphants. Since SEC apoptosis precedes neutrophil dispersal, we propose that apoptotic cell debris acts as danger-associated molecular patterns (DAMPs), recruiting neutrophils from the CHT. Apoptosis is a tightly regulated developmental process essential for removing excess cells, maintaining tissue homeostasis, and shaping organ morphology (Elmore, 2007). However, dysregulated apoptosis in the skin is related to several skin diseases, such as toxic epidermal necrolysis, psoriasis, and skin cancer (Raj et al., 2006). In zebrafish, neutrophil mobilization to the skin has been observed in mutants or morphants lacking functional *tnfr2*, clathrin interactor 1a (*clint1a*), or serine peptidase inhibitor Kunitz type 1a (*spint1a*, also known as *hai1a*), with some of these genes associated with keratinocyte hyperproliferation (Candel et al., 2014; Dodd et al., 2009; Mathias et al., 2007). Noticeably, only Clint1a deficiency has been shown to cause both increased apoptosis and proliferation of keratinocytes. These zebrafish models share key characteristics with human psoriasis, including keratinocyte hyperproliferation, chronic inflammation, excessive immune cell infiltration, and epithelial-mesenchymal transition-like features in keratinocytes (Martinez-Navarro et al., 2019). Although the mechanisms underlying psoriasis remain unclear, our Lpar1-deficient model may serve as an additional platform for studying this disease. Further investigation of keratinocyte proliferation, morphology, and tight junction integrity in Lpar1-deficient zebrafish could shed light on LPAR1’s role in epidermal homeostasis and broaden our understanding of inflammatory skin disorders.

### Lpar1 regulates neutrophil retention through *cxcl12a* expression in the CHT

In addition to apoptosis in the skin, we found that *cxcl12a* expression in the CHT was downregulated in the Lpar1 morphants. This downregulation occurred before neutrophils began to disperse, suggesting that reduced *cxcl12a* expression makes neutrophils more prone to mobilize in response to increased apoptosis in the skin.

Paredes-Zuniga et al. (2017) did not observe the neutrophil dispersal phenotype in both Cxcl12a and its receptor Cxcr4b loss-of-function mutants at 3 dpf, but they observed more circulating neutrophils in the bloodstream in both mutants at 7 dpf. Additionally, they found increased recruitment of neutrophils to tail fin transection sites, supporting our idea that the reduction of Cxcl12a would make neutrophils more mobilized. Isles et al. (2019) further verified the role of Cxcl12a-Cxcr4b signaling in neutrophil recruitment by using CRISPR interference (CRISPRi) to knock down Cxcl12a and Cxcr4b and the Cxcr4 antagonist AMD3100 to block the signaling. They again found increased neutrophil recruitment in Cxcr4 crispants but not in Cxcl12a crispants, possibly due to a reduction in total neutrophil numbers. Taken together, previous findings support our hypothesis that reduced *cxcl12a* expression augments neutrophil release from the CHT toward tissue damage, such as apoptotic SECs.

LPA-LPAR1 signaling has been implicated in the regulation of CXCL12-CXCR4 signaling. In a murine vascular injury model, inhibition of LPAR1 and LPAR3 using the inhibitor Ki6425 reduced CXCL12 expression in neointimal tissue, and incubating carotid vessels with unsaturated LPAs induced CXCL12 expression (Subramanian et al., 2010). Similarly, Wang et al. (2014) demonstrated that treatment of ovarian cancer cells with LPA induced CXCL12 secretion in a dose-dependent manner. These studies suggest a potential role for LPA-LPAR1 signaling in regulating CXCL12 secretion.

Regarding the transcriptional regulation of CXCL12, the activation of the phosphatidylinositol-3 kinase (PI3K)/Akt/specificity protein-1 (Sp1) signaling pathway has been shown to upregulate CXCL12 gene expression in CXCL12-abundant reticular (CAR) cells of the bone marrow (Adapala et al., 2019). Consistent with this regulatory link, we previously reported similar expression patterns of *lpar1* and *cxcl12a* in the CHT during early development (Lee et al., 2008). Furthermore, recent single-cell transcriptomic analyses identified high *lpar1* expression within the *cxcl12a*-rich stromal cell population in zebrafish (Sur et al., 2023). These findings collectively support a model in which Lpar1 positively regulates *cxcl12a* expression, thereby modulating neutrophil retention and mobilization during early hematopoietic development.

In addition to its role in regulating *cxcl12a* expression in the CHT, we also examined whether Lpar1 influences *cxcr4b* expression in neutrophils. Previous studies have shown that Lpar1 is expressed at low levels in human neutrophils under steady-state conditions, but is significantly upregulated in patients with pneumonia (Rahaman et al., 2006). In mice, Lpar1 expression is minimal in neutrophils recruited to the lung following bleomycin-induced injury; however, functional expression has been demonstrated, as Lpar1-deficient neutrophils fail to respond to LPA stimulation in vitro (Tager et al., 2008; Miyabe et al., 2019). These findings suggest that Lpar1 can be expressed in neutrophils across species, yet its expression in zebrafish neutrophils has not been previously characterized. Here, we report for the first time that Lpar1 is expressed in zebrafish neutrophils at steady state at 3 dpf. Despite this expression, loss of Lpar1 does not affect *cxcr4b* levels at this stage. These results indicate that Lpar1 regulates neutrophil retention in the CHT primarily through modulation of *cxcl12a* expression, rather than via direct regulation of *cxcr4b* in neutrophils.

## Conclusions

Our study reveals a novel role for Lpar1 in regulating neutrophil distribution in zebrafish. We observed that the absence of Lpar1 disrupts neutrophil dynamics by increasing apoptosis in SECs, impairing the retention signal for neutrophils in the CHT, and ultimately leading to their mobilization to the skin. These findings enhance our understanding of Lpar1’s function in neutrophil dynamics during steady state and early developmental stages. Therefore, investigating the role of Lpar1 in the development of certain skin diseases associated with immunodeficiency may provide insights into the underlying mechanisms.

## ACKNOWLEDGEMENTS

We thank the Taiwan Zebrafish core facility for providing fish lines and Ms. Yi-Chun Chuang in Technology Commons in National Taiwan University for excellent technical assistance with confocal microscopy. The work was supported by the National Science and Technology Council, Taiwan (NSTC-111-2311-B-002 -019 -MY3) to SJL.

