## Supplementary material for "Stay or Stray: Lpar1 regulates neutrophil retention and epidermal homeostasis in early zebrafish development": Suppl figures S1-6

### SUPPLEMENTARY FIGURES

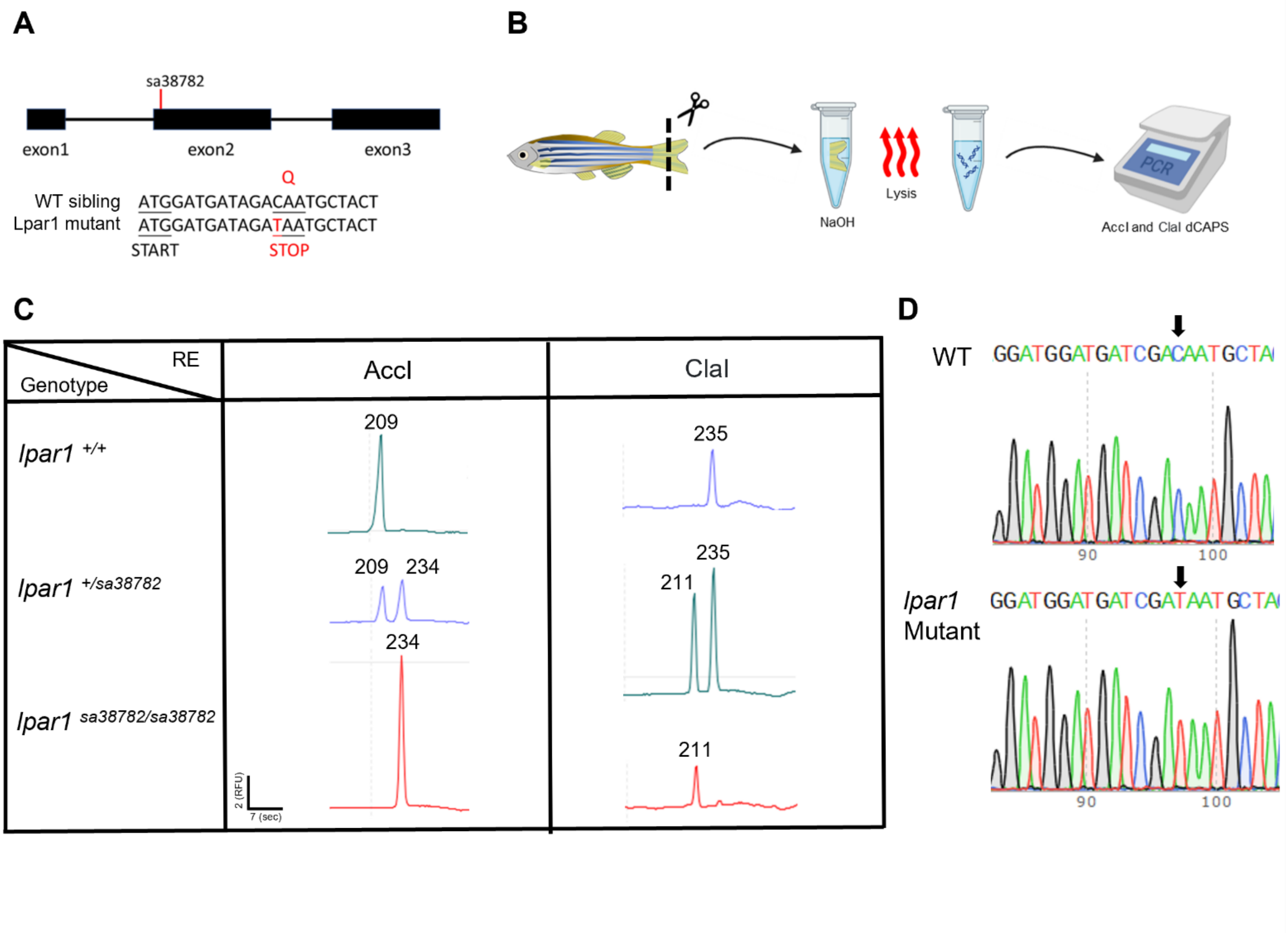


**Figure S1.** *Genotyping the Lpar1 sa38782 mutant.*

(A) The sa38782 mutant allele has a point mutation in the *lpar1* exon 2, which is a C to T mutation that leads to a premature start codon near the translation initiation site (START). As a result, a truncated Lpar1 protein with only four amino acids is predicted to be formed. (B) Workflow for genomic DNA extraction from adult zebrafish tail fin tissue for genotyping. Caudal fin fragments were dissected and placed into 0.2 mL PCR tubes containing 100 µL of 50 mM NaOH. Samples were incubated at 95 °C for 30 minutes to lyse the tissue and release genomic DNA. Subsequently, 10 µL of 1 M Tris-HCl (pH 8.0) was added to neutralize the solution. Due to the absence of an accessible restriction enzyme cut site near the mutation site of *lpar1*, we utilized the Derived Cleaved Amplified Polymorphic Sequences (dCAPS) assay for genotyping the mutant via restriction endonuclease (RE) digestion using AccI and ClaI enzymes. (C) Restriction fragment length polymorphism (RFLP) analysis was performed by capillary electrophoresis. For the AccI-based dCAPS assay, the 234 bp PCR product is cut in the wild-type allele, yielding a 209 bp fragment (the 25 bp fragment is not detectable), whereas the mutant remains uncut. For the ClaI assay, the enzyme cleaves only the mutant allele, producing a

211 bp fragment from the 235 bp amplicon; the 24 bp fragment is not detectable. The scale bar represents migration time in sec (X-axis), and fluorescence intensity is shown in relative fluorescence units (RFU, Y-axis). (D) Sanger sequencing confirmed the C-to-T point mutation in the mutant allele and verified the absence of additional mutations in the amplicons. Arrows indicate the position of the wild-type cytosine (C) and mutant thymine (T) nucleotides.

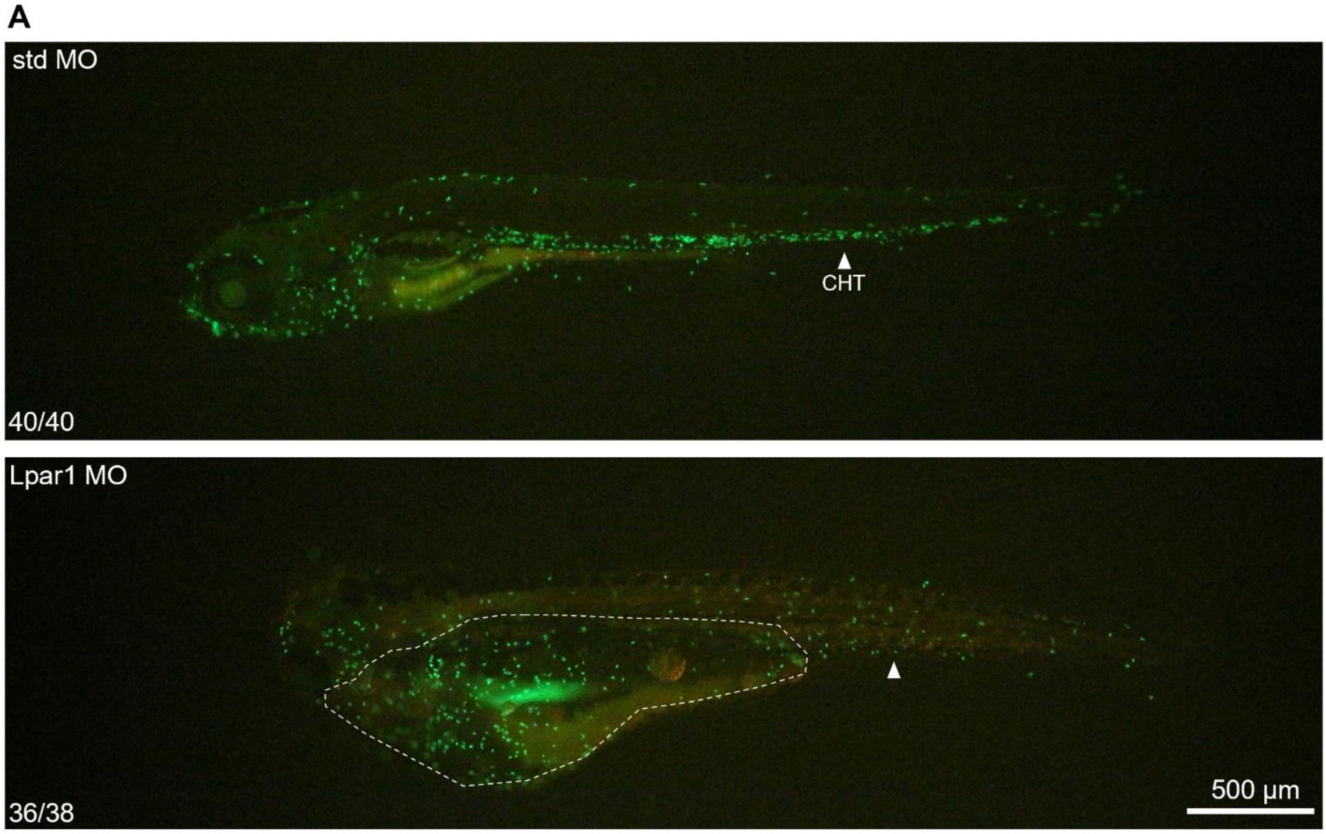


**Figure S2.** *Neutrophil accumulates at the edema site in Lpar1 morphant.*

**(A)** Representative images of *Tg(mpx:EGFP)* zebrafish embryos injected at the one-cell stage with 1.25 ng of either standard control MO (std MO) or Lpar1 splicing-blocking MO (Lpar1 MO). Images were captured at 5 days post-fertilization (dpf) under bright-field and fluorescence using epifluorescence microscopy. At 5 dpf, Lpar1 morphants exhibited pericardial and trunk edema. EGFP-labeled neutrophils were observed accumulating in the edema regions (white dashed circles), with apparent dispersal from the caudal hematopoietic tissue (CHT, white arrowheads), indicating altered neutrophil distribution in Lpar1 morphants.

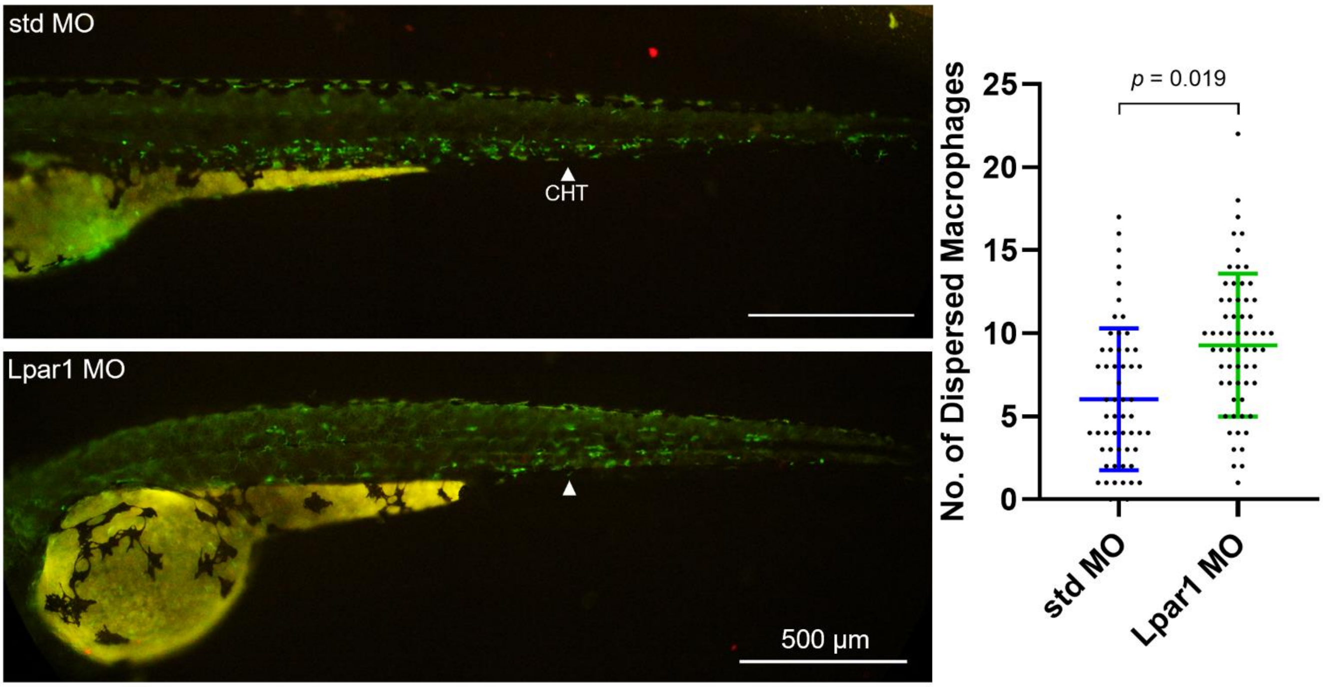


**Figure S3.** *Knockdown of Lpar1 causes the dispersal of macrophages.*

Representative images of *Tg(mpeg1:EGFP)* zebrafish embryos injected at the one-cell stage with 1.25 ng of either standard control MO (std) or *Lpar1* MO oligonucleotides (Lpar1), imaged under bright-field and epifluorescence microscopy at 3 days post-fertilization (dpf). Macrophages expressing EGFP are visible in green. Dispersal of macrophages from the caudal hematopoietic tissue (CHT) is indicated by white arrows. Quantification of dispersed macrophages within the defined region (as described in Fig. 1) was performed on 7–20 larvae per group across four independent experiments. Data are presented as mean ± s.d. Statistical analysis was conducted using Student's unpaired *t*-test.

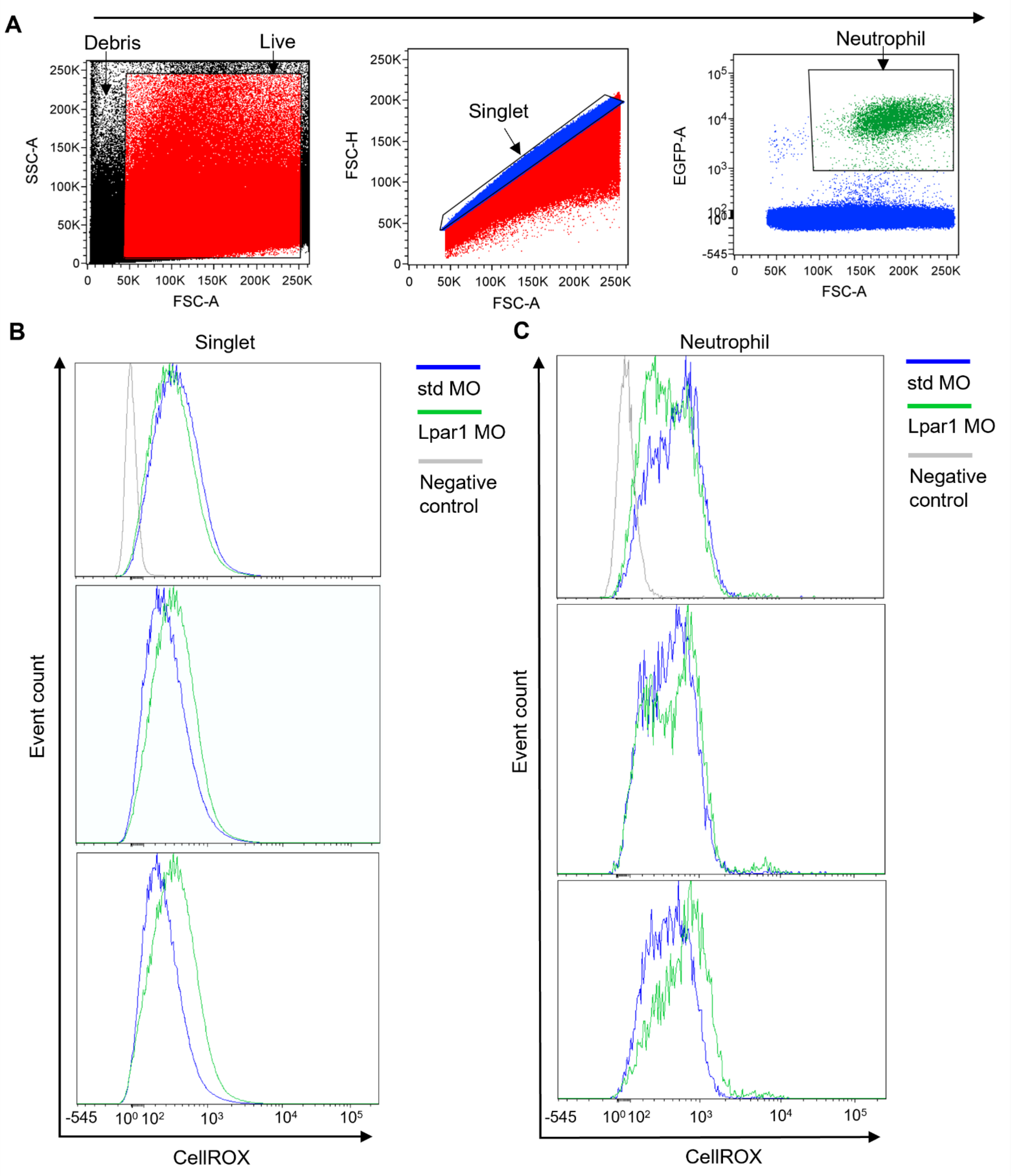


**Figure S4.** *Flow cytometry gating strategy for quantifying ROS levels in whole larvae and neutrophils of 3-dpf zebrafish.*

(A) Representative gating strategy for identifying neutrophils. Forward scatter (FSC) and side scatter (SSC) density plots were used to distinguish cells based on size and granularity. Viable cells were gated (live) by excluding debris. Singlet cells (singlet) were subsequently gated using FSC-H vs. FSC-A plots to eliminate doublets. Within the viable singlet population, neutrophils were identified based on EGFP fluorescence intensity and size. A distinct cluster with high EGFP intensity was defined as the neutrophil population. (B, C) Embryos were injected at the one-cell stage with 1.25 ng of either standard control MO (std MO) or Lpar1 splice-blocking MO (Lpar1 MO). Reactive oxygen species (ROS) levels were assessed by CellROX fluorescence intensity. Unstained controls (gray) served as negative controls. Histograms represent data from the same biological replicates shown in Fig. 6B, C (N = 3). (B) ROS levels in whole larvae were measured in the singlet population defined in (A): std MO (purple), Lpar1 MO (yellow), and negative control (gray). (C) ROS levels in neutrophils were measured in the EGFP$^+$ neutrophil population defined in (A); std MO (red), Lpar1 MO (blue), and negative control (gray).

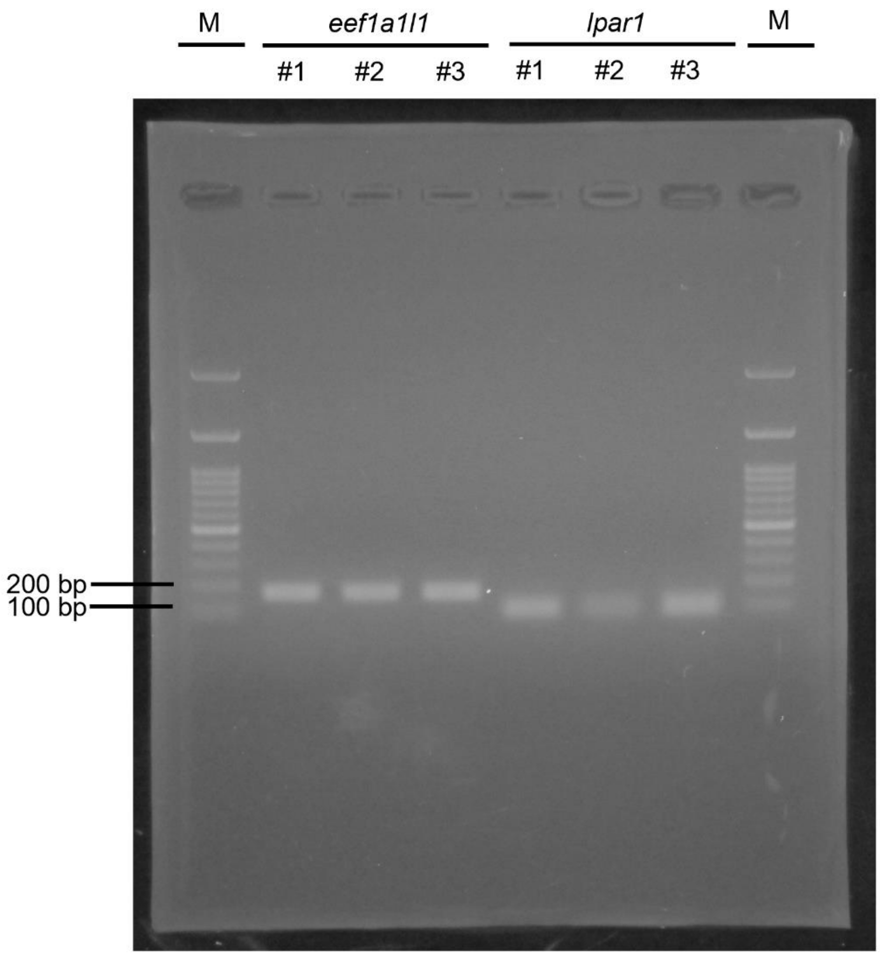


**Figure S5.** Lpar1 is expressed in neutrophils of 3 dpf zebrafish larvae.
RT-PCR gel electrophoresis shows a clear band corresponding to the *lpar1* amplicon (91 bp), amplified from cDNA derived from the standard control (std MO) group used in Fig. 6B. Eef1a1l1 (167 bp) serves as a positive control. Data are shown for three biological replicates (#1–3). M indicates the DNA ladder.

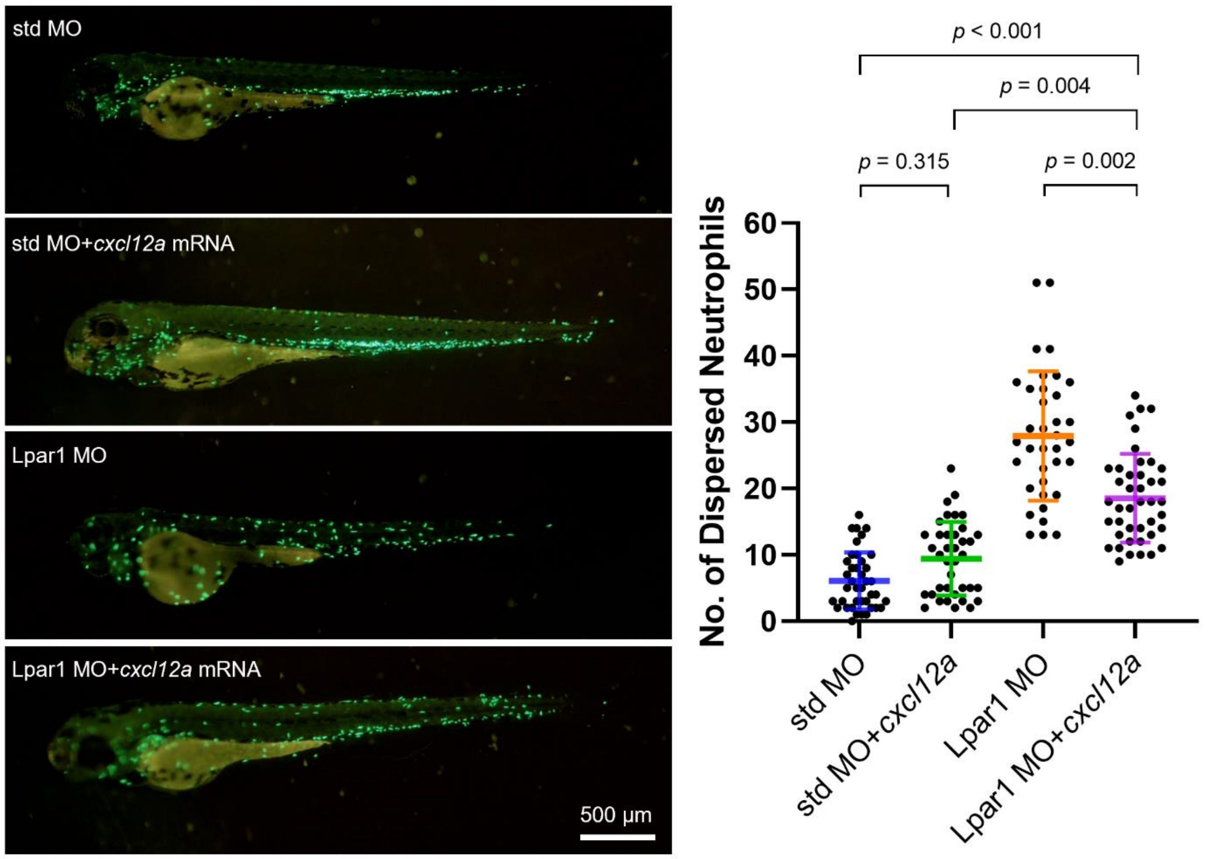


**Figure S6.** *Ectopic expression of cxcl12a partially rescues the neutrophil dispersal in Lpar1 morphants. Tg(mpx:EGFP)* zebrafish embryos were injected at the one-cell stage with 1.25 ng of either standard control MO (std), *Lpar1* splicing-blocking MO (Lpar1), or *Lpar1* MO co-injected with 200 pg of *cxcl12a* mRNA. Images were captured at 3 days post-fertilization (dpf) under bright-field and epifluorescence microscopy. Dispersed neutrophils were quantified using 11–15 larvae per group across three independent experiments. Data are presented as mean ± s.d., and analyzed using one-way ANOVA followed by Tukey's post hoc test.
